# Proinflammatory cytokines mediate pancreatic β-cell specific alterations to Golgi morphology via iNOS-dependent mitochondrial inhibition

**DOI:** 10.1101/2025.01.29.635550

**Authors:** Sandra E. Blom, Riley M. Behan-Bush, James A. Ankrum, Ling Yang, Samuel B. Stephens

**Affiliations:** Fraternal Order of Eagles Diabetes Research Center, University of Iowa, Iowa City, IA, USA; Department of Internal Medicine, Division of Endocrinology and Metabolism, University of Iowa, Iowa City, IA, USA; Roy J. Carver Department of Biomedical Engineering, University of Iowa, Iowa City, IA, USA; Department of Anatomy and Cell Biology, University of Iowa, Iowa City, IA, USA

**Author notes:** Lead author: Samuel B. Stephens, Ph.D., Fraternal Order of Eagles Diabetes Research Center, Departments of Internal Medicine, Division of Endocrinology and Metabolism, Department of Anatomy and Cell Biology, University of Iowa, Iowa City, IA 52246, USA.

**Keywords:** Type 1 diabetes, β-cells, Golgi, cytokines, nitric oxide, iNOS, Nos2, mitochondria, metabolism, glycosylation

## Abstract

Type 1 diabetes (T1D) is caused by the selective autoimmune ablation of pancreatic β-cells. Emerging evidence reveals β-cell secretory dysfunction arises early in T1D development and may contribute to diseases etiology; however, the underlying mechanisms are not well understood. Our data reveal that proinflammatory cytokines elicit a complex change in the β-cell’s Golgi structure and function. The structural modifications include Golgi compaction and loss of the inter-connecting ribbon resulting in Golgi fragmentation. Our data demonstrate that iNOS generated nitric oxide (NO) is necessary and sufficient for β-cell Golgi re-structuring. Moreover, the unique sensitivity of the β-cell to NO-dependent mitochondrial inhibition results in β-cell specific Golgi alterations that are absent in other cell types, including α-cells. Collectively, our studies provide critical clues as to how β-cell secretory functions are specifically impacted by cytokines and NO that may contribute to the development of β-cell autoantigens relevant to T1D.

## Introduction

Type 1 Diabetes (T1D) results from the autoimmune destruction of insulin-producing pancreatic β-cells^1^. Recent studies have defined an asymptomatic period that occurs years before clinical onset where β-cell mass declines and defects in β-cell secretory function emerge^2^. While the cellular mechanisms contributing to β-cell dysfunction during this asymptomatic period are not fully understood, proinflammatory cytokines, including IL-1β, IFN-γ, and TNF-α released from activated immune cells are thought to play a key role in disease development^3^. Following IL-1β and IFN-γ mediated upregulation of inducible nitric oxide synthase (iNOS), production of micromolar levels of nitric oxide (NO) mediate several inhibitory and destructive actions in β-cells^4–11^. These include suppression of mitochondrial oxidative metabolism, inhibition of insulin secretion, ER stress, DNA damage, and cell death. Although cytokines and NO are associated with β-cell damage, the effects are reversible for up to 36 hours^12^. Moreover, cytokines and NO can also induce the expression of antiviral and antimicrobial genes that may initially afford β-cells protection against DNA damage-induced apoptosis and viral replication^13–18^. These latter observations indicate that cytokine signaling in β-cells plays complex and potentially paradoxical roles, which may initially trigger protective mechanisms, but long term, induce cellular damage and death^19^. Taken together, these findings underscore the need to further investigate β-cell responses to inflammatory signals.

Early in T1D development, defects in β-cell secretory function may contribute to disease progression by facilitating or augmenting immune activation^2,20^. These defects include impaired insulin maturation^21,22^, mis-trafficking of secretory proteins^23,24^, formation of hybrid peptide neoantigens^25–27^, and increased release of extracellular vesicles and exosomes^28–31^. Although strong evidence supports a role for ER stress in mediating some of these secretory defects^8,32^, the impact of proinflammatory cytokines on the Golgi apparatus are not well understood^33,34^. Importantly, over a third of proteins encoded by the human genome traverse the Golgi enroute to the cell surface for insertion into the plasma membrane, secretion, or delivery to the endolysosomal system. Thus, defects in Golgi functions could significantly contribute to protein mis-sorting and dysregulated protein processing already identified in β-cells prior to T1D onset^34^. In addition, the polarized organization of the Golgi from *cis-* to *trans-* provides spatial separation of glycosyl transferases for stepwise assembly of protein glycans. Of note, the Golgi contains over 200 glycosyl transferases and is the primary organelle for glycan addition to cell surface proteins. Dysregulated protein glycosylation has been documented in human diseases that coincide with altered Golgi structure^37–41^ but have yet to be reported in T1D. Because protein glycosylation can be a critical determinant in antigen recognition and immunogenicity^42–44^, defects in β-cell protein glycosylation could contribute to the development of neoantigenic signals that stimulate immune activation.

In this report, we examined Golgi structure and function following proinflammatory cytokine exposure in rat, mouse, and human β-cells. Using confocal and electron microscopy, we identified striking alterations in Golgi morphology that include Golgi compaction, shortening of the Golgi cisternae, and loss of the inter-connecting ribbon leading to Golgi fragmentation. We demonstrate that Golgi structural changes coincide with decreased expression of cell surface glycoproteins. Importantly, these alterations were independent of changes in Golgi protein expression or cell death. Using *Nos2* (iNOS) KO mice, we show that alterations in Golgi structure are NO dependent. Additionally, we show that cytokine/NO mediated Golgi remodeling is due to metabolic suppression in β-cells. Comparisons of NO sensitivity between β-cells and non-β-cells revealed that Golgi alterations may be unique to β-cells due to their reliance on aerobic metabolism for energy homeostasis. Taken together, our data highlight significant alterations to Golgi structure in early T1D models that may provide clues to the defects in secretory functions and the specificity of β-cell autoimmune effects within the islet.

## Materials and methods

### Cell culture and reagents

INS1 832/3 cells (a gift from Dr. Christopher Newgard) were cultured as previously described^45^. HEK293A cells (Thermo Life Technologies) were cultured in DMEM supplemented with 10 % fetal bovine serum, and 1 % penicillin and streptomycin. αTC1 cells (a gift from Dr. Yumi Imai) were cultured in low-glucose (5.5 mM) DMEM supplemented with 10 % fetal bovine serum, 0.02% bovine serum albumin, 0.01 mM non-essential amino acids, and 1 % penicillin and streptomycin. Cell culture reagents were from Thermo Life Technologies unless specified otherwise. Recombinant mouse and human cytokines, IL-1β, TNF-α, and IFN-γ, were from Fisher Scientific (R&D). (Z)-1-[N-(3-aminopropyl)-N-(3-ammoniopropyl) amino]diazen-1-ium-1,2-diolate (DPTA) was purchased from VWR. Transfection of GFP-reporter plasmids was achieved using Lipofectamine 3000 (Thermo Life Technologies). Mouse islets were isolated via collagenase V digestion and purified using Histopaque 1077 and 1119 (Sigma). Mouse islets were cultured in RPMI supplemented with 10 % fetal bovine serum and 1 % penicillin and streptomycin and maintained at 37°C in 5 % CO_2_. For introduction of adenoviral reporters, cells or islets were transduced with ∼1-5 × 10^7^ IFU/mL or ∼ 1-5 × 10^8^ IFU/mL adenovirus for 18 h and assayed 72-96 h post-treatment. Human islets obtained from Prodo Laboratories and Alberta Diabetes Institute IsletCore were cultured in RPMI (Sigma) supplemented with glucose to 5.5 mM, 10% fetal bovine serum and 1% penicillin and streptomycin. All donor families gave informed consent for the use of pancreatic tissue in research. Donor information can be found in Supplemental Table 1.

### Animal studies

Wild-type (WT) C57BL/6J mice and whole-body *Nos2* (iNOS) KO mice were obtained from Jackson Laboratories and bred in-house. Mouse studies equally represent male and female mice. All animal protocols were approved by the University of Iowa Institutional Animal Use and Care Committee.

### Plasmids and viruses

GFP-GRASP65 plasmid^46^ was a gift from Dr. Yanzhuang Wang (University of Michigan). Rat insulin promoter (RIP)-driven eGFP was assembled into a modified pAd-PL/DEST via multi-site Gateway cloning using LR Clonase II plus^47^. AdRIP-RFP virus was purchased from the University of Iowa Viral Vector Core. Recombinant adenoviruses were amplified in HEK293 cells and purified by cesium chloride gradient. All sequences were verified by the Iowa Institute of Human Genetics, University of Iowa.

### Glucose-stimulated insulin secretion

For static measures of insulin secretion, INS1 832/3 cells or pools of 10 islets were incubated in secretion assay buffer^45^ containing 2.5 mM glucose for 1 h at 37° C followed by incubation at 12 mM (cells) or 16.7 mM (mouse islets) glucose for 1 h. Islets or INS1 832/3 cells were lysed in radioimmunoprecipitation assay buffer for content determination. Insulin (secreted and content) was measured by ELISA (rodent 80-INSMR-CH10; ALPCO).

### Cellular respiration

For measurements of cellular respiration, GLPBIO Cell Counting Kit 8 (CCK8) protocol was followed. INS1 832/3 cells were seeded at a density of ∼5000 cells/well. Cytokine treatment was added 24 h post cell seeding. CCK8 reagent was added following 18 h cytokine treatment.

### Flow Cytometry

INS1 832/3 cells were lifted in PBS and pelleted at 500 x g’s for 5 min before propidium iodide (PI) staining for 10 min (Invitrogen). The number of PI positive cells was measured using a Cytek Northern Lights spectral cytometer. Before collecting experimental data, the cytometer was calibrated using SpectroFlo Cytometer QC Beads (Cytek). Instrument voltage settings were determined using an unstained and positive PI-stained INS1 832/3 cell population. The experimental data was assessed using the SpectroFlo software (Cytek). Cells were gated using singlet discrimination (SSC-A/SSC-H and FSC-A/FSC-H). Then from the singlet parent population, the percentage of PI-positive cells was reported.

### Immunoblot Analysis

Clarified cell lysates were resuspended in LDS sample buffer (Thermo Life Technologies), resolved on 4-12 % NuPAGE gels (Thermo Life Technologies), and transferred to supported nitrocellulose membranes (BioRad). Membranes were probed with diluted antibodies raised against GM130 (mouse, BD Transduction, 610823, 1:1000), Golgin97 (rabbit, Proteintech, 12640-1-AP, 1:1000), or γ-tubulin (mouse, Sigma, T5326). Donkey anti-rabbit and anti-mouse antibodies coupled to IR-dye 680 or 800 (LI-COR) were used to detect primary antibodies. Blots were developed using an Odyssey CLx Licor Instrument.

### Quantitative RT-PCR

RNA from INS1 832/3 cells was harvested using a Zymo RNA minikit. RNA from mouse and human islets was harvested using the RNeasy Microkit (QIAGEN). cDNA was synthesized in an iScript reaction (BioRad). Real-time PCR was performed using either ABI 7700 sequence detection and software (Applied Biosystems) or QuantStudio-7 PRO sequence detection system and software Design & Analysis.

### Fluorescence Microscopy and Imaging

For imaging studies, isolated islets were gently dispersed into monolayer sheets using Accutase (Sigma-Aldrich) as previously described^48^. INS1 832/3 cells, and mouse islet monolayers were plated on HTB9-coated coverslips (Electron Microscopy Services) and cultured overnight. Human islets were plated onto Collagen IV (Thermo Life Sciences) coated coverslips and cultured overnight^49^.

For immunostaining, cells and islets were fixed in 10% neutral-buffered formalin. Permeabilized cells were incubated overnight with antibodies raised against GM130 (mouse, BD Transduction 610,823), Golgin97 (rabbit, Proteintech 12640-1-AP), insulin (guinea pig, Fitzgerald 70R-10659), or TGN38 (mouse, Novus Biologicals NB300-575) as indicated. Highly cross-adsorbed fluorescent conjugated secondary antibodies (whole IgG, donkey anti-mouse AlexaFluor 647, donkey anti-rabbit Cy3, and donkey anti-guinea pig AlexaFluor488; Jackson ImmunoResearch) were used for detection. Cells were counterstained with DAPI (Sigma) and mounted using Fluorosave (Calbiochem). Images were captured on a Leica SP8 confocal microscope using a HC PL APO CS2 63x/1.40 oil objective with 3x zoom as z-stacks (5 per set, 0.33 μm step, 0.88 μm optical section) and deconvolved (Huygen’s Professional). Golgi fragment number and Golgi area were determined using a surface rendering module in Imaris (Bitplane) from GM130 immunostaining and normalized per cell number. Primary β-cells were identified by insulin positive staining to ensure β-cell specific measurements.

For glycan imaging studies, islets were infected with AdRIP-GFP or AdRIP-RFP to mark β-cells and fixed in 10% neutral-buffered formalin. Fixed islets were incubated with fluorescently conjugated lectins Wheat-germ agglutinin (WGA; Thermo Life Sciences) or Concanavalin A (ConA; Thermo Life Sciences) for 10 min. Cells were counterstained with DAPI (Sigma) and mounted using Fluorosave (Calbiochem). Images were captured on a Leica SP8 confocal microscope using a HC PL APO CS2 63x/1.40 oil objective with 3x zoom as z-stacks (5 per set, 0.33 μm step, 0.88 μm optical section). ImageJ was used to measure lectin fluorescence intensity within GFP/RFP positive cells and intensity normalized by cell number.

### Ultrastructure

All EM related reagents were from Electron Microscopy Sciences. Isolated islets were fixed in 2.5% glutaraldehyde, 4% formaldehyde cacodylate buffer (72-96 h) at 4⁰ C. Tissue was post-fixed in fresh 1% OsO_4_ for 1 h, dehydrated using a graded alcohol series followed by propylene oxide and embedded in Epon resin. Resin blocks were trimmed with a glass knife, cut to ultrathin (50-70 nm) sections with a diamond knife, and mounted on Formvar-coated copper grids. Grids were double contrasted with 2% uranyl acetate then with lead citrate. Images were captured at 10,000x magnification by a Hitachi HT7800 transmission electron microscope. β-cells were identified by the presence of morphologically distinct dense core secretory granules. Golgi cisternal length and width was determined using the line segment tool in Fiji NIH software. Individual Golgi stacks were counted and defined as dilated if greater than 50% of counted cisternae were > 50 nm in width^50^.

### Statistical Analysis

Data are presented as the mean ± S.D. For statistical significance determinations, data were analyzed by the two-tailed unpaired, Student’s t test or by ANOVA with post-hoc analysis for multiple group comparisons as indicated (GraphPad Prism). A *p*-value < 0.05 was considered significant

## Results

### Proinflammatory cytokines elicit Golgi fragmentation in INS 832/3 cells

Proinflammatory cytokines contribute to β-cell dysfunction during the asymptomatic period of T1D development^3^. While β-cell secretory defects have been linked to ER stress^8,32^, alterations to other critical secretory organelles, such as the Golgi are less well-defined^33,34^. To address this, we examined INS1 832/3 cells treated with a cytokine cocktail containing IL-1β (0.01 ng/mL or 0.05 ng/mL)^17,51,52^, TNF-α (750 U/mL)^53–55^, and IFN-γ (750 U/mL)^53,54^. Following 18 h treatment, we observed a dose-dependent decrease in glucose-stimulated insulin secretion (GSIS, Fig. 1A), insulin content (Fig. 1B), and cellular respiration (Fig. 1C), consistent with previous reports. Next, we investigated the effect of proinflammatory cytokines on Golgi morphology. In control cells, we observed a continuous perinuclear Golgi ribbon network identified by *cis* (GM130) and *trans* (Golgin97) Golgi markers (Fig. 1D). In contrast, cytokine-treated INS1 832/3 cells contained numerous Golgi fragments. (Fig. 1D-E). Similar observations were made using GRASP65 (*cis*) and TGN38 (*trans*) markers (Fig. S1A-C). Golgi fragmentation was not accompanied by changes in protein or mRNA expression of *cis-* (GM130 or GRASP55), or *trans-* (TGN38 or Golgin97) Golgi markers (Figs. 1F-H, S1D). Taken together, these data identify unique alterations to β-cell Golgi structure in response to cytokines, independent of changes in Golgi protein expression.

**Figure 1.**
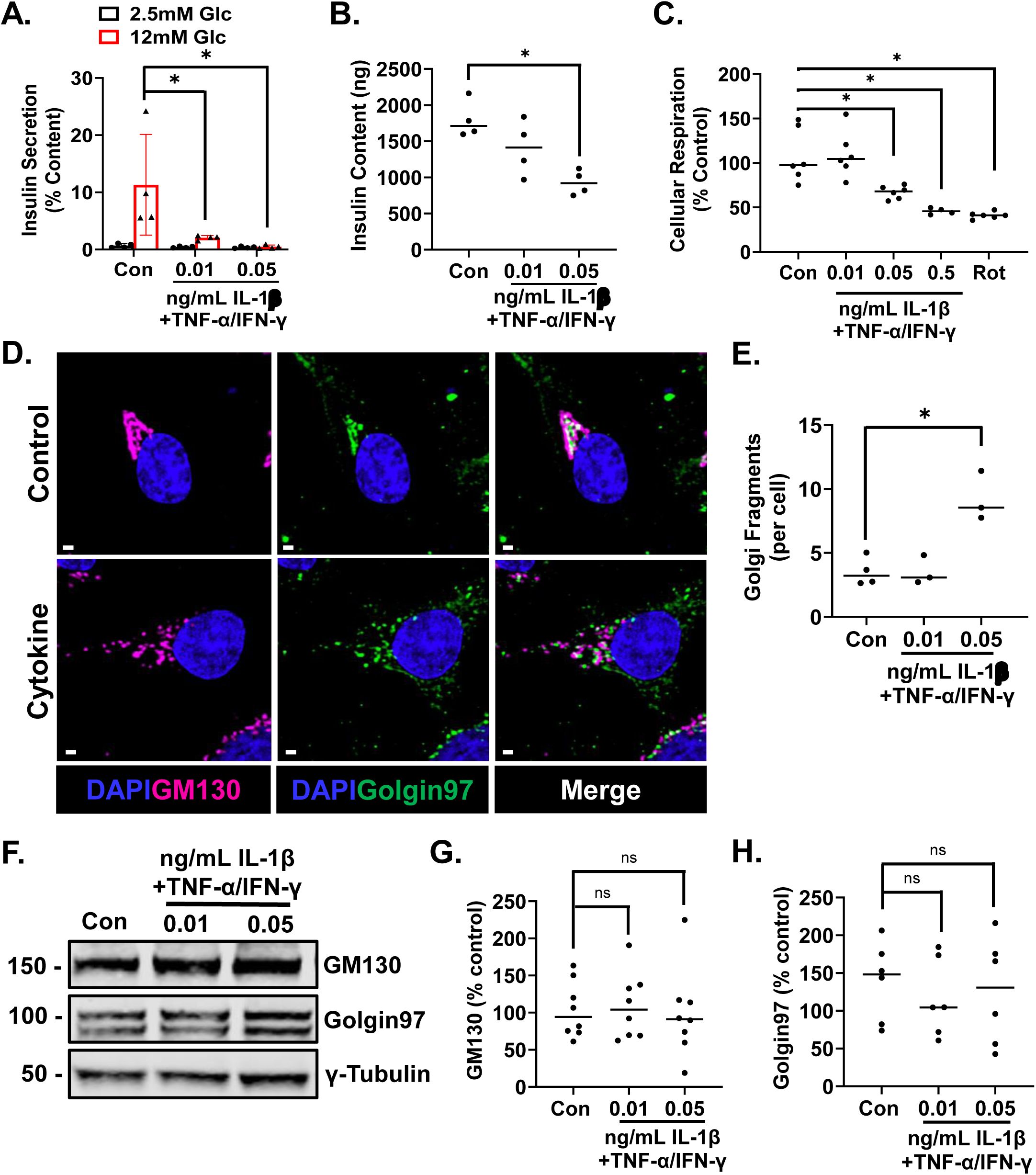
Proinflammatory cytokines drive Golgi fragmentation in INS1 832/3 cells. INS1 832/3 cells were untreated (control) or treated with cytokine cocktail (IL-1β, 0.01 ng/mL or 0.05 ng/mL; TNF-α, 750 U/mL; IFN-γ, 750 U/mL) for 18 h. (**A**) Glucose-stimulated insulin secretion was measured by sequential static incubation in media containing 2.5 mM Glc or 12 mM Glc for 1 h each as indicated. Data are normalized to insulin content. (**B**) Insulin content was determined from whole-cell lysates. (**C**) Cellular respiration was measured by MTT. Cells were treated with rotenone (1µM) for 15 min as a positive control. Data are normalized relative to untreated controls. (**D**-**E**) Cells were immunostained for GM130 (magenta) and Golgin97 (green), and counterstained with DAPI (blue). (**D**) Representative images are shown. Scale bar = 1 μm. (**E**) Golgi fragments were quantified from GM130 immunostaining and normalized per cell. (**F**) Whole cell lysates were analyzed by immunoblot (**F**) and normalized to γ-tubulin (**G**-**H**). (**A**-**C**, **E**, **G**-**H**) Individual experiments are presented as data points with the mean ± S.D. as indicated. *p < 0.05 by 2 way-ANOVA with repeated measures (**A**) or 1-way ANOVA with repeated measures (**B**-**C**, **E**, **G**-**H**).

### Proinflammatory cytokines drive Golgi compaction in primary β-cells

We next explored if Golgi structural changes were also evident in primary β-cells treated with cytokines. To determine this, isolated mouse islets were cultured with a proinflammatory cytokine cocktail containing IL-1β (0.02 ng/mL or 0.1 ng/mL)^17,51,52,55^, TNF-α (750 U/mL)^53–55^, and IFN-γ (750 U/mL)^53,54^. As expected, 18 h cytokine treatment resulted in a dose-dependent decrease in GSIS (Fig. 2A) and insulin content (Fig. 2B). Using these conditions, we investigated Golgi morphology. In untreated (control) mouse β-cells, we observed a continuous perinuclear Golgi ribbon network with distinct *cis*-(GM130) and *trans*-(Golgin97) Golgi compartments (Fig. 2C). Following cytokine treatment of mouse β-cells, we observed a striking compaction of the Golgi ribbon (Fig. 2C), rather than fragmentation detected in the INS1 832/3 model (Fig 1D-E). Quantitation confirmed a significant decrease in cytokine-treated β-cell Golgi area measured by GM130 marker staining (Fig. 2D). No significant difference in cell size (Fig. 2E) or Golgi protein expression (Fig. S2A-C) was detected.

**Figure 2.**
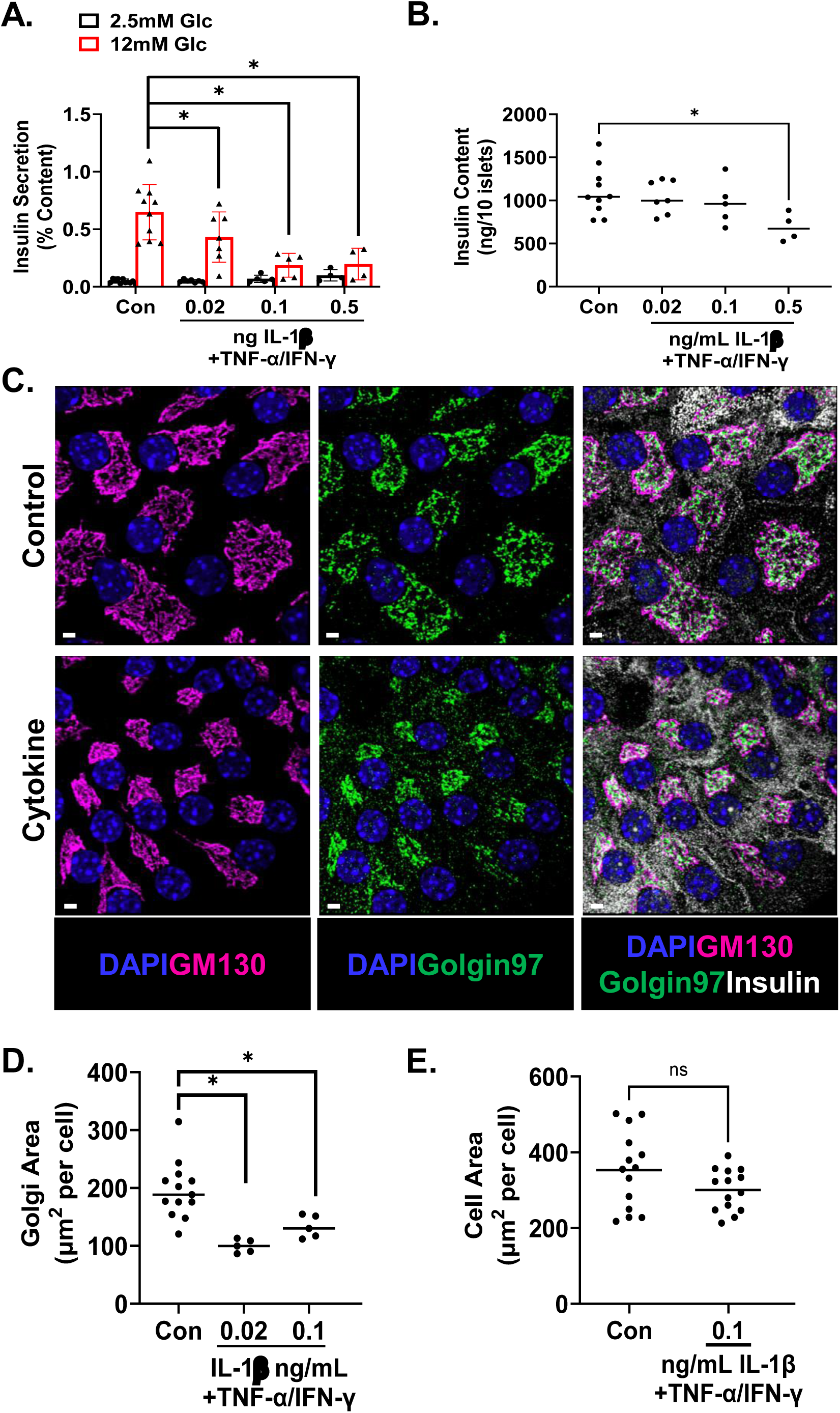
Proinflammatory cytokines induce Golgi compaction in primary mouse β-cells. Mouse islets were untreated (control) or treated with cytokine cocktail (IL-1β, 0.02 ng/mL or 0.1 ng/mL; TNF-α, 750 U/mL; IFN-γ, 750 U/mL) for 18 h. (**A**) Glucose-stimulated insulin secretion was measured by sequential static incubation in media containing 2.5 mM Glc or 16.7 mM Glc for 1 h each as indicated. Data are normalized to insulin content. (**B**) Insulin content was determined from whole-cell lysates. (**C**-**E**) Cells were immunostained for GM130 (magenta), Golgin97 (green), and insulin (white), and counterstained with DAPI (blue). (**C**) Representative images are shown. Scale bar = 3 μm. (**D**) Golgi area was quantified from GM130 staining of insulin positive cells and normalized per β-cell. (**E**) β-cell area was quantified from insulin staining and normalized per cell. (**A**-**B**, **D**-**E**) Individual mice are presented as data points with the mean ± S.D. as indicated. *p < 0.05 by 2 way-ANOVA with repeated measures (**A**) or 1-way ANOVA with repeated measures (**B**, **D**, **E**).

After observing Golgi structural changes in both INS1 832/3 and mouse primary β-cells, we explored Golgi morphology in human primary β-cells following cytokine exposure. To examine this, we utilized islets harvested from non-diabetic human donors. Islets were cultured with a cytokine cocktail containing IL-1β (0.5 ng/mL or 2.5 ng/mL)^53,54,56^, TNF-α (1500 U/mL)^56^, and IFN-γ (750 U/mL)^53,54^ for 48 h., which resulted in decreased insulin content (Fig. S3A). Using this cytokine cocktail, we investigated Golgi morphology. Similar to primary mouse β-cells (Fig. 2C-D), cytokine treatment resulted in substantial compaction of the Golgi (Fig 3A, B) without Golgi fragmentation (Fig 3C). We further investigated the loss of Golgi area in human primary β-cells by transmission electron microscopy (TEM). β-cells were readily identifiable by their morphologically distinct dense core secretory granules. Ultrastructural analysis of untreated controls revealed Golgi cisternae that were organized into long, thin stacks (Fig. 3D) averaging 1916 nm in length (Fig. 3E) and 256.9 nm in width (Fig. 3F). In contrast, cytokine treatment resulted in disrupted Golgi cisternae (Fig. 3D) that were significantly shorter (Fig. 3E). No difference in stack width (Fig. 3F) or stack number (Fig. 3G) was observed between control and cytokine treated human β-cells. Additionally, no significant difference in stack dilation (Fig. S3B) or vesiculation (Fig. S3C) was detected despite the disorganized appearance of the Golgi stacks (Fig. 3D). Collectively, these data identify profound, cytokine-mediated alterations in Golgi structure in three separate β-cell models.

**Figure 3.**
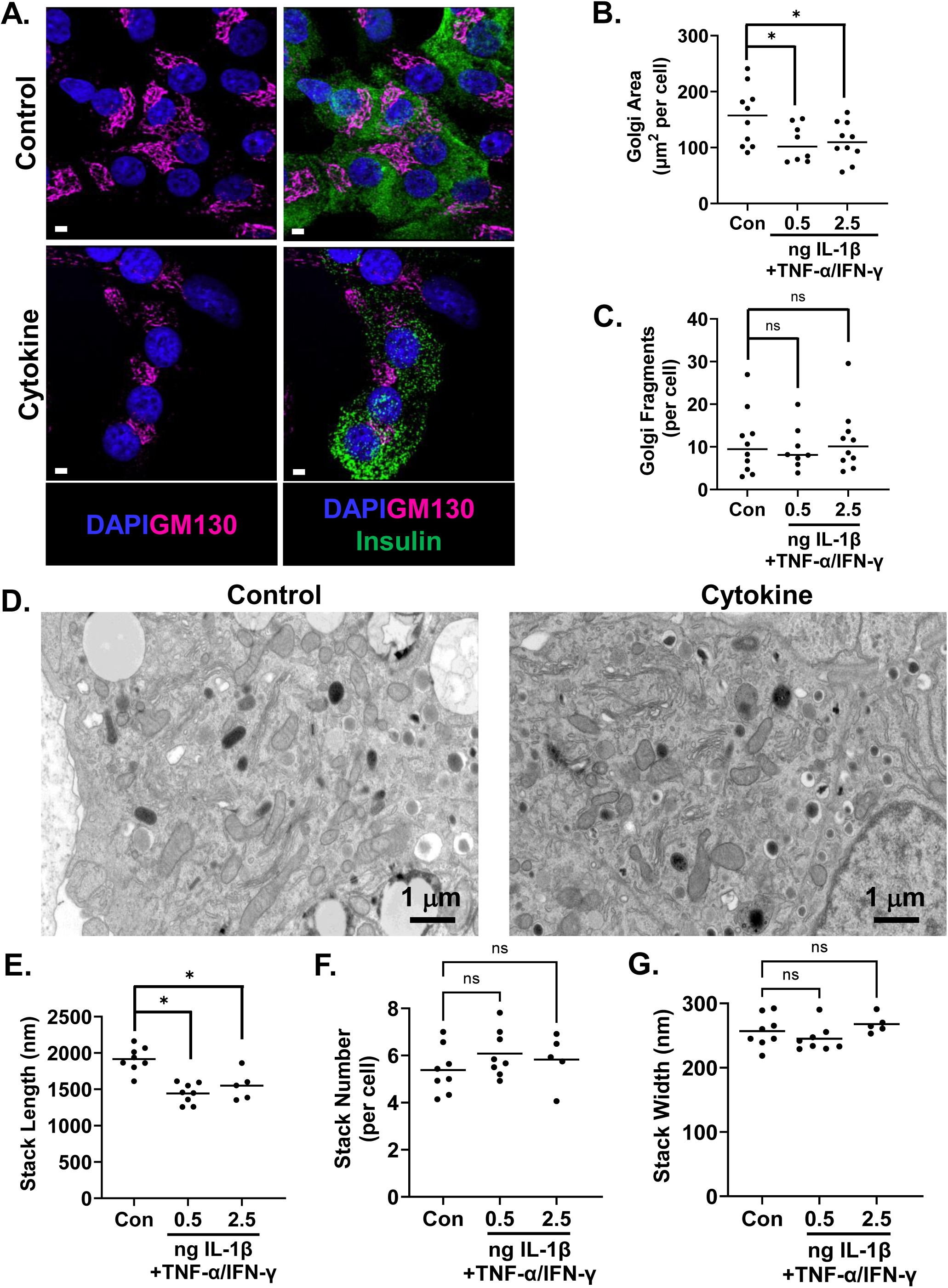
Proinflammatory cytokines elicit Golgi cisternal compaction in primary human β-cells. Non-diabetic human islets were untreated (control) or treated with cytokine cocktail (IL-1β, 0.5 ng/mL or 2.5 ng/mL; TNF-α, 1000 U/mL; IFN-γ, 750 U/mL) for 48 h. (**A**-**C**) Cells were immunostained for GM130 (magenta) and insulin (green) and counterstained with DAPI (blue). (**A**) Representative images are shown Scale bar = 3 μm. Golgi area (**B**) and Golgi fragments (**C**) were quantified from GM130 staining per insulin positive β-cell. (**D**-**G**) β-cells were examined by transmission electron microscopy. (**D**) Representative micrographs are shown. Golgi were analyzed (15 images/donor) for cisternal length (**E**), stacks per cell (**F**), and cisternal width (**G**). (**B**-**C**, **E-G**) Individual donors are presented as data points with the mean ± S.D. as indicated. * p < 0.05 by Student t test.

### Cytokine-induced Golgi remodeling is reversible

The effects of proinflammatory cytokines on metabolism and cell viability are reversible for up to 36 h^12^. To examine if Golgi alterations are also reversible, INS1 832/3 cells were treated with cytokine mix for 18 h followed by a 24 h or 48 h washout in normal growth media and compared to untreated and cytokine treated INS1 cells. As shown previously (Fig. 1D), 18 h cytokine treatment resulted in robust Golgi fragmentation as identified by GM130 staining (Fig. 4A, B). Following cytokine withdrawal and return to normal growth media, Golgi fragmentation was remediated within 24 h and stable for up to 48 h (recovery; Fig. 4A, B). Propidium iodide (PI) staining showed no significant increase in PI positivity following 18 h cytokine treatment or 48 h recovery (Fig. 4C). Similarly, no change in nuclear size of cytokine treated INS1 832/3 or mouse islet cells was detected (Fig. 4D, E). Taken together, these data demonstrate that β-cell Golgi alterations in response to proinflammatory cytokine treatment are fully reversible and independent of cell death.

**Figure 4.**
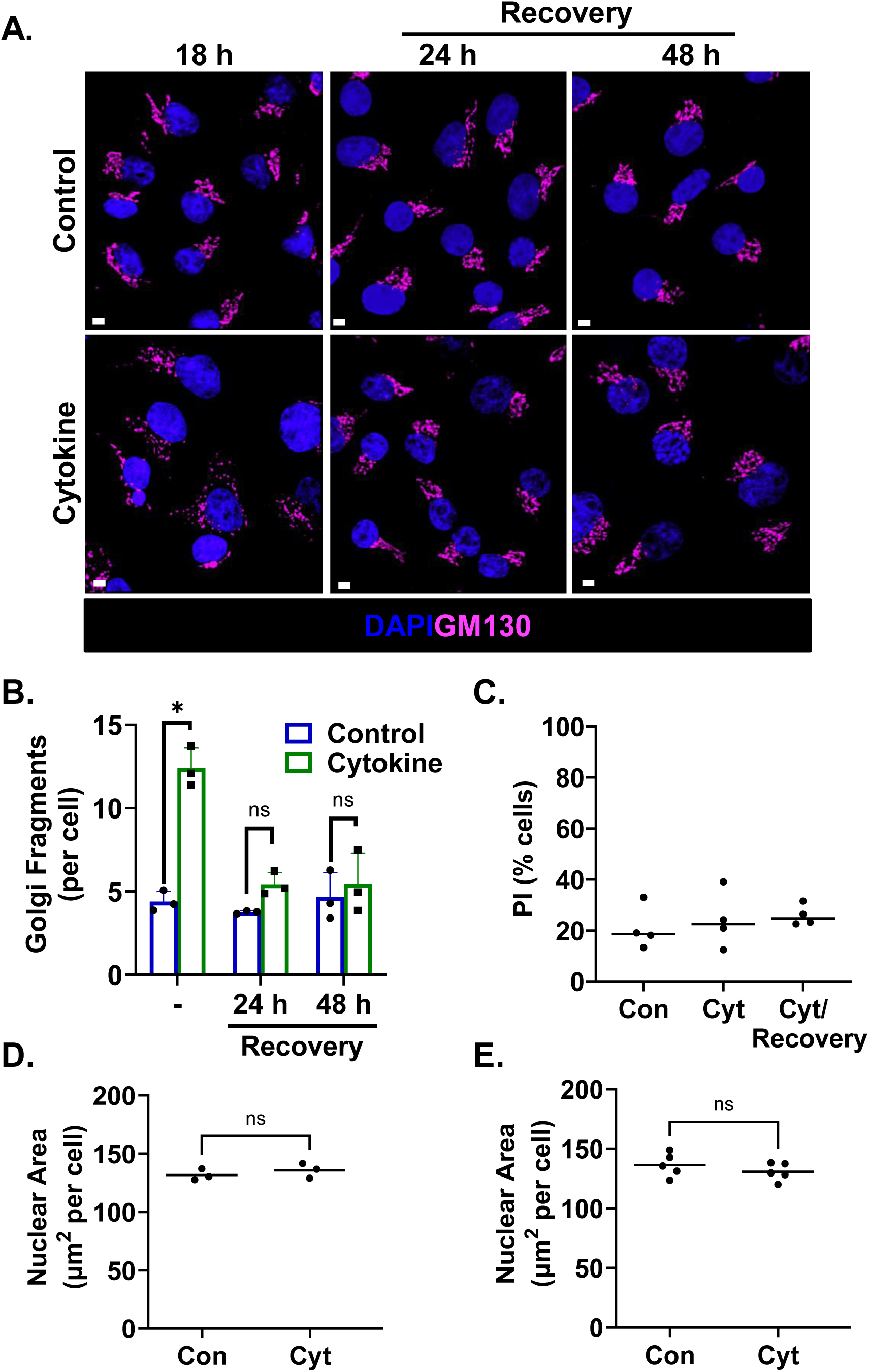
Cytokine-mediated Golgi fragmentation is reversible. INS1 832/3 cells were untreated (control) or treated with cytokine cocktail (IL-1β, 0.05 ng/mL; TNF-α, 750 U/mL; IFN-γ, 750 U/mL) for 18 h and returned to normal growth media for 24 h or 48 h (recovery) as indicated. (**A**-**B**) Cells were immunostained for GM130 (magenta) and counterstained with DAPI (blue). (**A**) Representative images are shown. Scale bar = 3 μm. (**B**) Golgi fragments were quantified per cell by GM130 staining. (**C**) Cell death was quantified from propidium iodide (PI) staining. (**D**) Nuclear area was quantified per cell using DAPI staining. (**E**) Mouse islets were untreated or treated with cytokine cocktail (IL-1β, 0.1 ng/mL; TNF-α, 750 U/mL; IFN-γ, 750 U/mL) for 18 h. Nuclear area was quantified per cell using DAPI staining. (**B**-**E**) Individual experiments or mice are presented as data points with the mean ± S.D. as indicated. *p < 0.05 by 2 way-ANOVA with repeated measures (**B**), 1-way ANOVA with repeated measures (**C**), or Student’s t-test (**D**-**E**).

### Proinflammatory cytokines alter Golgi functions

To determine if Golgi function is also altered by cytokine treatment, we examined cell surface protein glycosylation. In these studies, cell surface glycoproteins were detected by lectin staining of non-permeabilized mouse islet cells. β-cells were identified by expression of a rat insulin promoter driven GFP (AdRIP-GFP; Fig. 5A, C) or RFP (AdRIP-RFP; Fig. 5D, E). Untreated control β-cells showed expression of cell surface glycoproteins using wheat germ agglutinin staining (WGA; Fig. 5A, B), which binds N-acetyl glucosamine (GlcNAc) and sialic acid containing glycans. In contrast, 18 h cytokine treatment resulted in a marked decrease in cell surface WGA staining (Fig. 5A, B). Note, that WGA staining of glycoproteins from whole cell lysates revealed no difference following cytokine treatment (Fig. 5C), suggesting a specific decrease in cell surface glycoprotein abundance rather a global reduction. Similarly, using concanavalin A (ConA), which binds mannose-containing glycans, we identified a marked reduction in cell surface glycoprotein expression in cytokine-treated mouse β-cells. These studies suggest that Golgi structural alterations coincide with Golgi functional defects.

**Figure 5.**
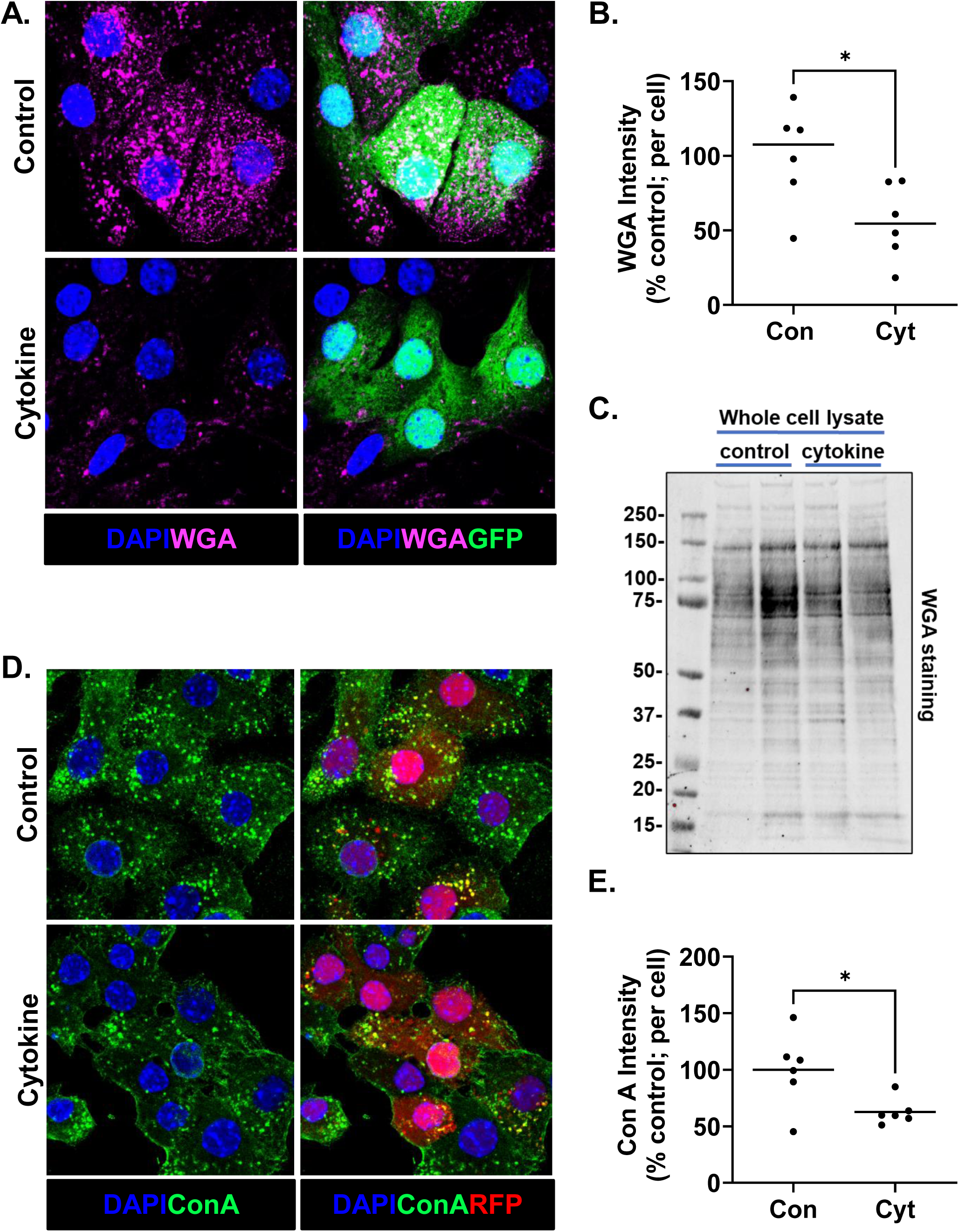
Proinflammatory cytokines decrease cell surface glycoproteins. Mouse islets were treated with AdRIP-GFP (**A**-**B**) or AdRIP-RFP (**D**-**E**) as indicated to identify β-cells. Mouse islets were untreated (control) or treated with cytokine cocktail (IL-1β, 0.1 ng/mL; TNF-α, 750 U/mL; IFN-γ, 750 U/mL) for 18 h. (**A**-**B**) Non-permeabilized cells were stained with WGA (magenta) and counterstained with DAPI (blue). (**B**) WGA fluorescence was quantified per cell and normalized to control cells. (**C**) INS1 832/3 cells were untreated (control) or treated with cytokine cocktail (IL-1β, 0.05 ng/mL; TNF-α, 750 U/mL; IFN-γ, 750 U/mL) for 18 h. Whole cell lysates were analyzed by WGA staining. (**D**-**E**) Non-permeabilized islet cells were stained with ConA (green) and counterstained with DAPI (blue). (**E**) ConA fluorescence was quantified per cell and normalized to control cells. (**B**, **E**) Individual mice are presented as data points with the mean. *p < 0.05 by Student’s t-test.

### iNOS is necessary and sufficient for cytokine-mediated Golgi alterations

IL-1β signaling in β-cells leads to upregulation of iNOS and micromolar production of NO, which accounts for the metabolic suppression of insulin secretion attributed to cytokine exposure^4–11^. Here, we sought to understand if iNOS and NO also contribute to cytokine-mediated alterations in Golgi morphology. Increased iNOS expression was confirmed following cytokine treatment in rat, mouse, and human models by RT-qPCR (Figure S4A-C). To explore if iNOS is necessary for Golgi compaction, islets from WT or *Nos*2 (iNOS) whole-body KO mice were treated with proinflammatory cytokine cocktail for 18 h. Mirroring our previous findings (Fig. 2C, D), WT β-cells displayed significant Golgi compaction compared to untreated controls (Fig. 6A, B). In contrast, Golgi area was unaffected by cytokines in *Nos2* KO β-cells with no observed compaction of the Golgi ribbon (Fig. 6A, B). This data suggests that iNOS is necessary for the cytokine-mediated Golgi remodeling to occur.

**Figure 6.**
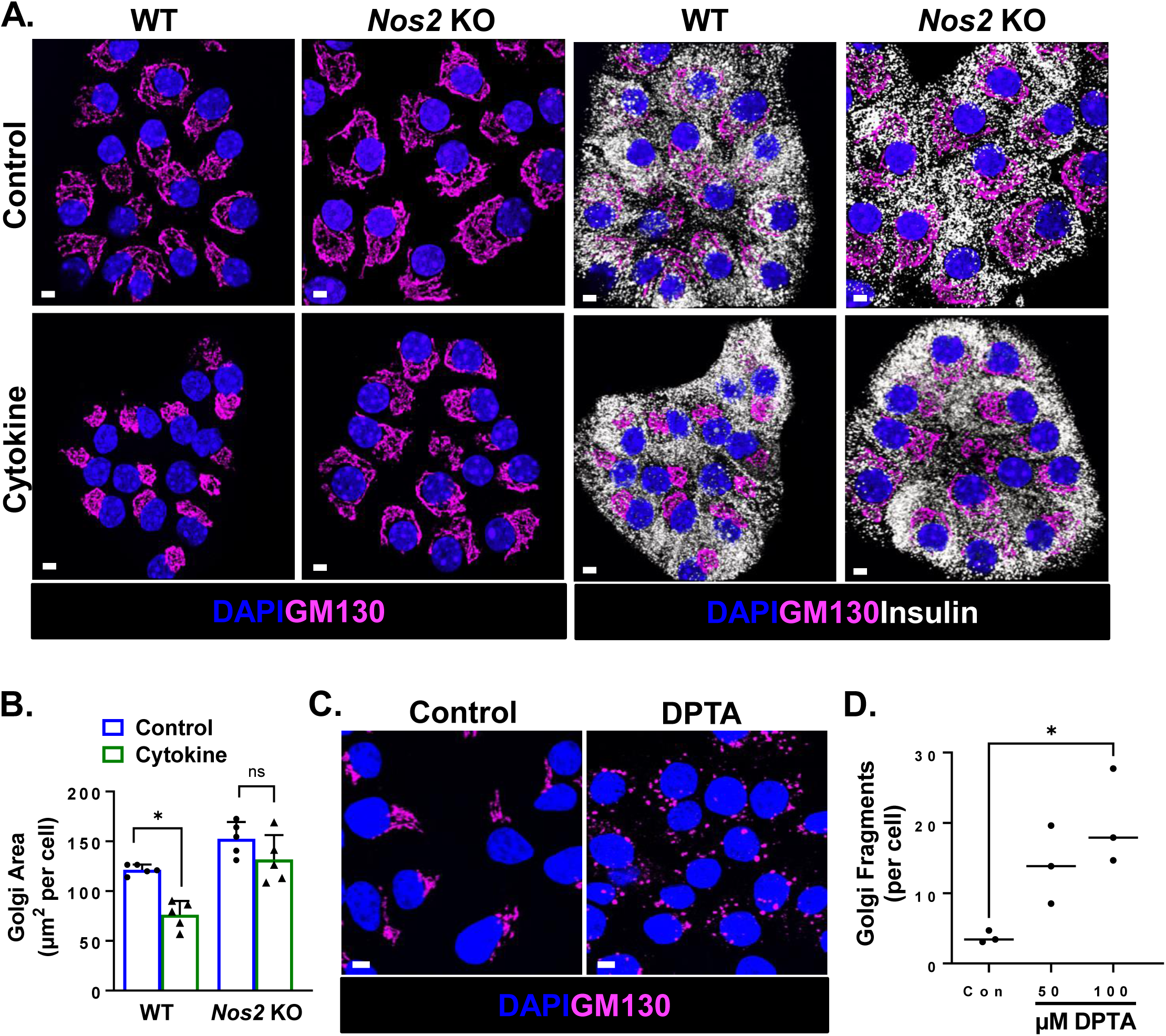
iNOS is necessary for cytokine-mediated Golgi remodeling. (**A**-**B**) Isolated islets from WT or *Nos2* KO mice were untreated (control) or treated with cytokine cocktail (IL-1β, 0.1 ng/mL; TNF-α, 750 U/mL; IFN-γ, 750 U/mL) for 18 h. (**A**-**B**) Cells were immunostained for GM130 (magenta) and insulin (white), and counterstained with DAPI (blue). (**A**) Representative images are shown. Scale bar = 3 μm. (**B**) Golgi area was quantified from GM130 staining per cell. (**C**-**D**) INS1 832/3 cells were untreated (control) or treated with DPTA (50 μM or 100 μM) for 4 h. Cells were immunostained for GM130 (magenta) and counterstained with DAPI (blue). (**C**) Representative images are shown. Scale bar = 5 μm. (**D**) Golgi fragments were quantified from GM130 staining per cell. (**B**, **D**) Individual mice or experiments are presented as data points with the mean ± S.D. as indicated. *p < 0.05 by 2 way-ANOVA with repeated measures (**B**) or 1-way ANOVA with repeated measures (**D**).

Because genetic deletion of iNOS prevented cytokine-mediated Golgi compaction, we next investigated whether NO was sufficient to alter Golgi structure. For this, we utilized the chemical NO donor, DPTA, which slowly decomposes to release micromolar levels of NO similar to iNOS^13^. Following 4 h DPTA treatment of INS1 832/3 cells, we observed a dose-dependent decrease in GSIS (Fig. S5A), mimicking our observations with cytokine exposure (Fig. 1A). We next investigated Golgi morphology following DPTA treatment. In control cells, Golgi structure presented as a continuous ribbon network (Fig. 6C). Contrasting this, in DPTA-treated cells, we observed robust Golgi fragmentation, mirroring cytokine treatment of INS1 832/3 cells (Fig. 6C, D). Taken together, these data show that iNOS and NO are necessary and sufficient for cytokine-mediated alterations of Golgi structure.

### Metabolic influence on Golgi structure

Recent studies have shown both α-cells and β-cells respond to cytokine exposure via iNOS induction^14,16–19^, yet β-cells are the primary target for autoimmune destruction in T1D and α-cell mass remains unchanged or increased^57^. To determine if cytokine signaling in α-cells results in Golgi alterations similar to β-cells, we treated αTC1 and INS1 832/3 cells with proinflammatory cytokine mix for 18 h. As expected, INS1 832/3 cells displayed robust Golgi fragmentation following cytokine treatment (Fig. 7A, B). In contrast, no change in Golgi structure was detected in cytokine treated αTC1 cells (Fig. 7A, B), despite strong upregulation of iNOS (*Nos2*) by cytokines in both cell types (Fig. 7C).

**Figure 7.**
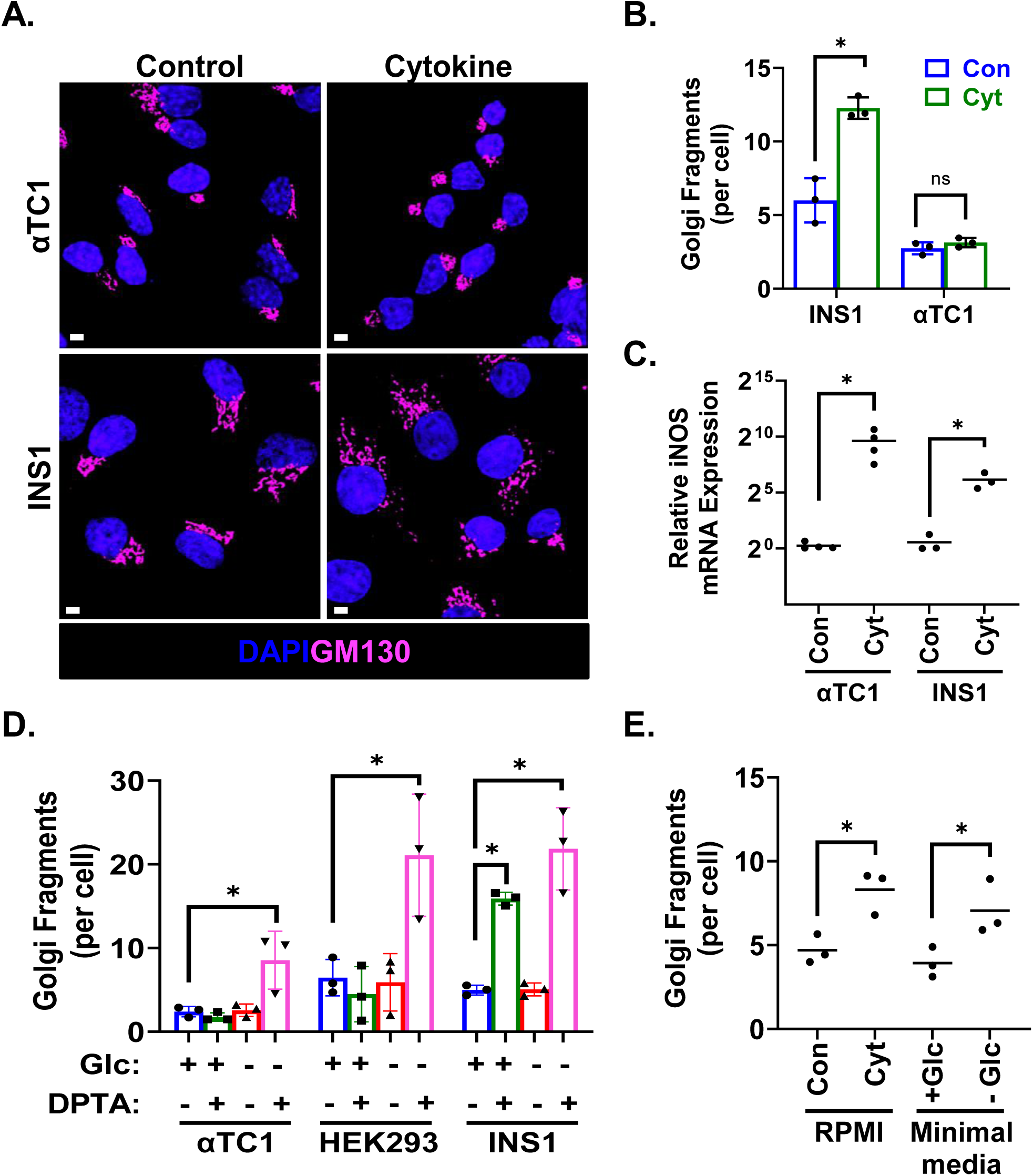
Metabolic differences reveal β-cell specific Golgi responses to NO. (**A**-**C**) αTC1 and INS1 832/3 cells were untreated or treated with cytokine cocktail (IL-1β, 0.05 ng/mL; TNF-α, 750 U/mL; IFN-γ, 750 U/mL) for 18 h. (**A**-**B**) Cells were immunostained for GM130 (magenta) and counterstained with DAPI (blue). (**A**) Representative images are shown. Scale bar = 3 μm. (**B**) Golgi fragments were quantified from GM130 staining per cell. (**C**) *Nos2* expression was quantified by RT-qPCR and normalized to untreated control cells. (**D**) αTC1, HEK293, and INS1 832/3 cells were cultured in growth media with or without glucose and untreated or treated with DPTA (100 μM) for 4 h as indicated. Cells were immunostained for GM130 and counterstained with DAPI. Golgi fragments were quantified from GM130 staining per cell. (**E**) INS1 832/3 cells were untreated (control) or treated with cytokine cocktail (IL-1β, 0.05 ng/mL; TNF-α, 750 U/mL; IFN-γ, 750 U/mL) for 18 h or cultured in minimal media with (+) or without (−) glucose (Glc, 12 mM) for 8 h. Cells were immunostained for GM130 and counterstained with DAPI. Golgi fragments were quantified from GM130 staining per cell. (**B**-**E**) Individual experiments are presented as data points with the mean ± S.D. as indicated. *p < 0.05 by 2-way ANOVA with repeated measures (**B**, **D**), Student’s t-test (**C**), or 1 way-ANOVA with repeated measures (**E**).

NO-mediated mitochondrial inhibition in β-cells results in robust suppression of ATP generating mechanisms^4–7,9,10^. In contrast, non-β-cells, including α-cells, can bypass NO suppression of mitochondrial function and maintain ATP production through anaerobic glycolysis^58,59^. Because cellular metabolism has been shown to regulate Golgi structure in various cell types^60,61^, we speculated that the distinct α-cell and β-cell Golgi responses to cytokines and NO was due to metabolic differences. Based on this idea, we explored the relationship between metabolic activity and Golgi structure in β-cells, α-cells, and non-islet cells using INS1 832/3, αTC1, and HEK293A cells, respectively. Cells were cultured in complete growth media supplemented with or without glucose and treated with the chemical NO donor, DPTA, for 4 h. Media lacking glucose had no effect on Golgi morphology in any of the three cell types tested (Fig. 7D), likely due to the presence of other fuels in the growth media, such as amino acids that feed directly into mitochondrial metabolism. While DPTA-treated INS1 832/3 cells displayed severe Golgi fragmentation (Figs. 7D, S6) as previously shown (Fig. 6C, D), no change in Golgi morphology was observed in αTC1 or HEK293A cells following DPTA treatment alone (Figs. 7D, S6). Instead, Golgi fragmentation was only detected in αTC1 and HEK293A cells upon DPTA treatment if glucose was also absent from the growth media (Figs. 7D, S6). To further explore the metabolic dependence for Golgi structure in β-cells, we next cultured INS1 832/3 cells in a minimal media for 8 h supplemented with or without glucose as the sole source of metabolic fuel. INS1 832/3 cells cultured in the absence of fuel (no glucose) exhibited robust Golgi fragmentation, whereas glucose supplementation maintained Golgi structure (Figs. 7E, S7A). Similarly, direct mitochondrial inhibition using the Complex I inhibitor, rotenone, also led to robust Golgi fragmentation (Fig. S7B, C). Collectively, these data demonstrate that Golgi structure is dependent on metabolic activity, and that non-β-cells, including α-cells, may be insensitive to cytokine and NO-mediated Golgi disruption provided that a glycolytic substrate is available.

## Discussion

T1D results from the selective autoimmune destruction of pancreatic β-cells^1^. Longitudinal studies of high-risk patients demonstrate that a prolonged asymptomatic period can precede clinical presentation with evidence of β-cell dysfunction up to 5 years prior to T1D diagnosis^2^. These observations suggest that slowly evolving changes between immune cell and β-cell interactions are critical to T1D development. A major hurdle in identifying disease modifying therapies is a limited understanding of the β-cell’s role in early disease progression^1,2,20^. Prior to symptomatic onset, inflammatory mediators, including proinflammatory cytokines, likely contribute to disease development by altering or impairing critical β-cell functions^4–11^. Cytokines are known to impair β-cell mitochondrial activity, elicit ER stress, and decrease β-cell viability. In the current study, we show that proinflammatory cytokines also elicit a complex change in the β-cell’s Golgi structure in human, mouse, and rat models. Cytokine treatment results in Golgi fragmentation in rat INS1 832/3 cells and compaction of the Golgi in primary mouse and human β-cells. While the spectrum of phenotypes between INS1 832/3 cells and primary β-cells is distinct, all three models display significant modifications to overall Golgi structure. We also show that Golgi structural changes are accompanied by significant decreases in β-cell surface glycoprotein abundance, which we speculate are due to changes in Golgi functions of protein glycosylation and trafficking. Finally, our data demonstrate that iNOS generated NO is necessary and sufficient for β-cell Golgi alterations that act via suppression of β-cell metabolism. Due to the β-cell’s strict reliance on mitochondrial function for ATP generation^58,59^, our data reveal that β-cells are uniquely sensitive to iNOS/NO effects as compared to α-cells. Based on these data, we propose that cytokines contribute to the early etiology of T1D development through Golgi-directed alterations that are specific to the pancreatic β-cell.

The Golgi is a critical organelle regulating the trafficking and post-translational modification of secreted and cell surface membrane proteins. Compromised Golgi integrity has been linked to human diseases of neurons, such as Huntington’s and Alzheimer’s diseases^62–65^, multiple cancer types^66^, and more recently by our group in T2D β-cells^67^, but is relatively unexplored in T1D^33,34,68^. Our data suggest that cytokine disruption of β-cell Golgi structure significantly alters glycoprotein abundance at the cell surface. While additional studies are needed to understand what further changes may occur at the β-cell surface, glycans are well-regarded as fundamental molecular signatures for discriminating between self and non-self antigens^69^. Indeed, the repertoire of glycan-binding proteins and lectins present on immune cells have been linked to a number of autoimmune diseases, such as rheumatoid arthritis, through changes in protein glycosylation and lipidation patterns^70,71^. Because dampening autoimmune signaling remains a critical strategy for attenuating or reversing T1D onset^1^, these observations highlight a critical need for further investigation into the relationship of Golgi dysfunction with the formation of autoantigens, particularly on the β-cell surface, and T1D development.

Evidence for Golgi dysfunction in T1D etiology has emerged from multiple reports of protein mis-sorting in β-cells. For example, elevated levels of circulating proinsulin compared to C-peptide are seen in both high risk autoantibody positive, yet asymptomatic, individuals, and in patients with long-standing disease^21^. These observations suggest that impaired proinsulin maturation and trafficking is a common defect in T1D. Secretion of ER-resident protein chaperones, such as BiP, from β-cells suggests a breakdown of the *cis*-Golgi localized KDEL receptor system necessary for retrieval and retrograde transport of ER proteins back to the ER^24,72^. Importantly, secreted ER chaperones have been shown to activate macrophages in co-culture that may contribute to loss-of-tolerance during immune surveillance of pancreatic β-cells ^28^. Defects in Golgi sorting are also seen with the mis-trafficking of endolysosomal proteins. The mis-sorting and/or retention of cathepsin D in the insulin secretory granule can catalyze the formation of hybrid peptide neoantigens^25–27^, which may elicit early immune activation. Thus, disrupting Golgi functions could provide a molecular switch for generating neoantigenic signals that modify how β-cells are surveilled by immune effectors early in T1D development.

IL-1β is extensively characterized to inhibit β-cell metabolic functions, specifically aerobic metabolism, through NO inactivation of iron-sulfur cluster-containing metabolic enzymes, such as aconitase^4–11^. Using *Nos2* (iNOS) KO mice, our work showed that iNOS/NO is necessary for changes in Golgi structure following cytokine treatment. Based on reports in other cell types that energy homeostasis is critical for maintaining Golgi structure^60,61^, we further evaluated if the loss of cellular metabolism could drive the reorganization of the Golgi in β-cells. Indeed, our analysis revealed that direct mitochondrial inhibition or simply removing fuel led to rapid loss of β-cell Golgi integrity. In addition, the reversibility of the Golgi response to cytokines is consistent with the restoration of cellular metabolism upon cytokine withdrawal^12^. Most cell types have the ability to switch from aerobic (mitochondrial) metabolism to anaerobic glycolysis for ATP production via LDH, which is necessary to recycle NAD^+^ for continued glycolysis^58,59^. Pancreatic β-cells do not express LDH, nor the monocarboxylate carrier, MCT-1^73^, and therefore are uniquely rigid in their reliance on aerobic metabolism for energy homeostasis. By comparing β-cells to α-cells and non-β-cells, our data show that Golgi alterations in response to NO may be unique to β-cells. Non-β-cells, including α-cells, can circumvent NO-mediated Golgi disruption, provided that a glycolytic substrate is available for anaerobic metabolism. In contrast, the metabolic inflexibility of the β-cell results in a unique susceptibility to Golgi modifications following cytokine and NO-mediated metabolic inactivation. Collectively, these observations reveal potential clues as to how β-cells are specifically impacted by cytokines, compared to other islet cell types, which could lead to the generation of β-cell autoantigens in the development of T1D.

## Resource Availability

This study did not generate any new unique materials.

## Acknowledgements

We would like to thank the donor families for pancreatic islet tissue. We also thank Dr. Chantal Allamargot for expert technical assistance. We acknowledge the use of the University of Iowa Central Microscopy Research Facility. This work was supported by Breakthrough T1D (SRA-2024-1553), the National Institutes of Health (R01 DK140093), the US Department of Defense (W81XWH-20-1-200), and a Fraternal Order of Eagles Diabetes Research Center Pilot and Feasibility Catalyst grant to S.B.S. and the National Institutes of Health (F31 DK139619) to S.E.B. S.B.S. is the guarantor of this work and, as such, had full access to all the data in the study and takes responsibility for the integrity of the data and the accuracy of the data analysis. All data are available upon reasonable request.

## Author contributions

S.E.B and S.B.S conceived and designed studies. S.E.B and S.B.S collected samples, performed the experiments, and analyzed the data. R.M.B and J.A.A provided flow cytometry reagents, and technical and analytical support. L.Y provided *Nos2* KO mice. S.E.B and S.B.S. wrote the manuscript.

## Supplemental Figure Legends

**Supplemental Figure 1.**
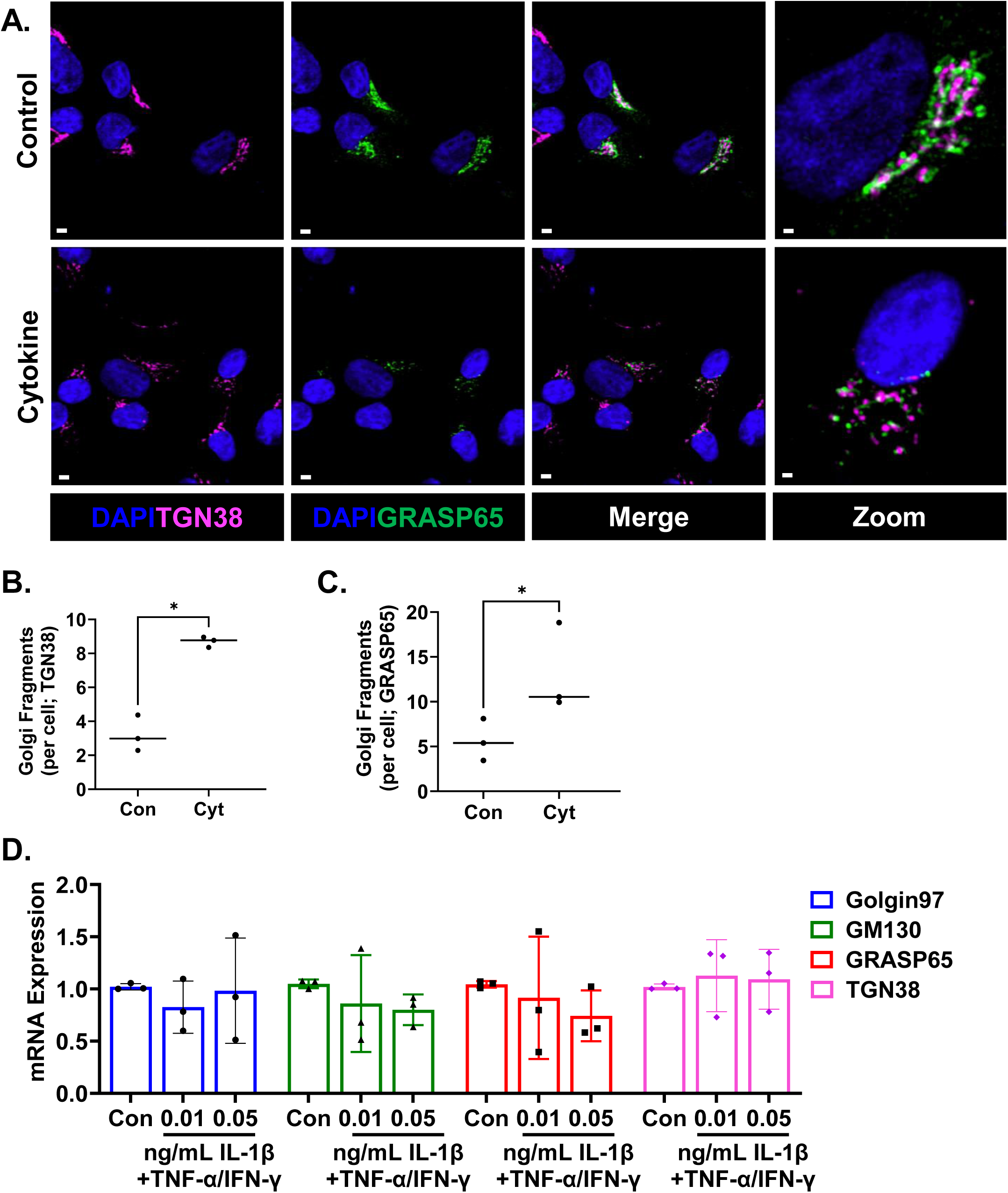
Golgi fragmentation in INS1 832/3 cells following cytokine treatement. INS1 832/3 cells were untreated or treated with cytokine cocktail (IL-1β, 0.05 ng/mL; TNF-α, 750 U/mL; IFN-γ, 750 U/mL) for 18 h. (**A**-**C**) Cells were transfected with GFP-GRASP65 (green), immunostained with TGN38 (magenta), and counterstained with DAPI (blue). (**A**) Representative images are shown. Scale bar = 3 μm. Zoomed imaged scale bar = 1 μm. Golgi fragments were identified by TGN38 (**B**) and GRASP65 (**C**) staining per cell. (**D**) mRNA expression of Golgi proteins was examined by qRT-PCR. (**B**-**D**) Individual experiments are presented as data points with the mean ± S.D. as indicated. *p < 0.05 by Student’s t-test (**B-C**) or 2-way ANOVA with repeated measures (**D**).

**Supplemental Figure 2.**
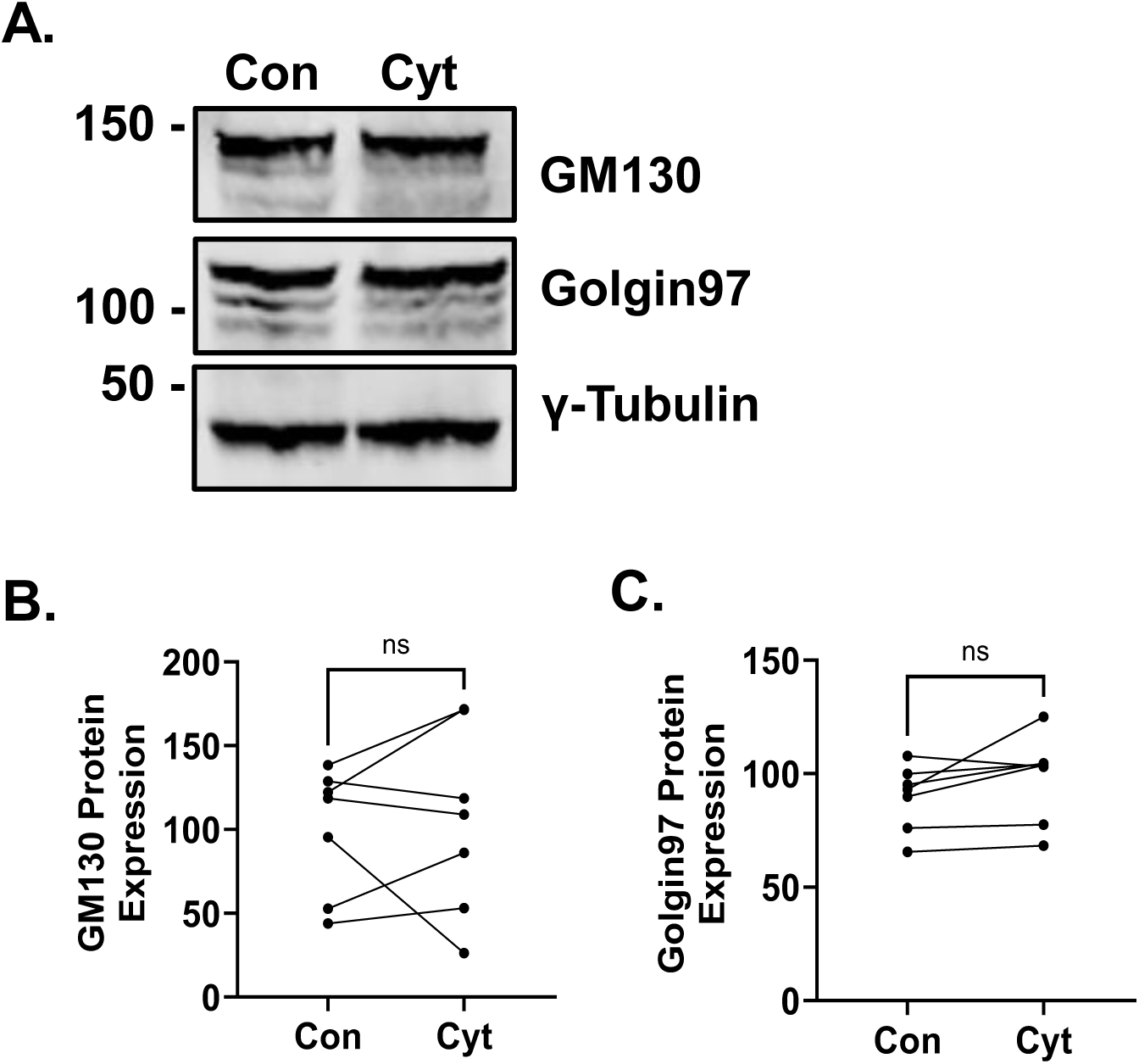
Mouse islet Golgi protein expression following cytokine treatment. Mouse islets were untreated or treated with cytokine cocktail (IL-1β, 0.1 ng/mL; TNF-α, 750 U/mL; IFN-γ, 750 U/mL) for 18 h. Whole cell lysates were analyzed by immunoblot (**A**) and normalized to γ-tubulin (**B**-**C**). (**B-C**) Paired data from individual mice are presented. *p < 0.05 by Student’s t-test.

**Supplemental Figure 3.**
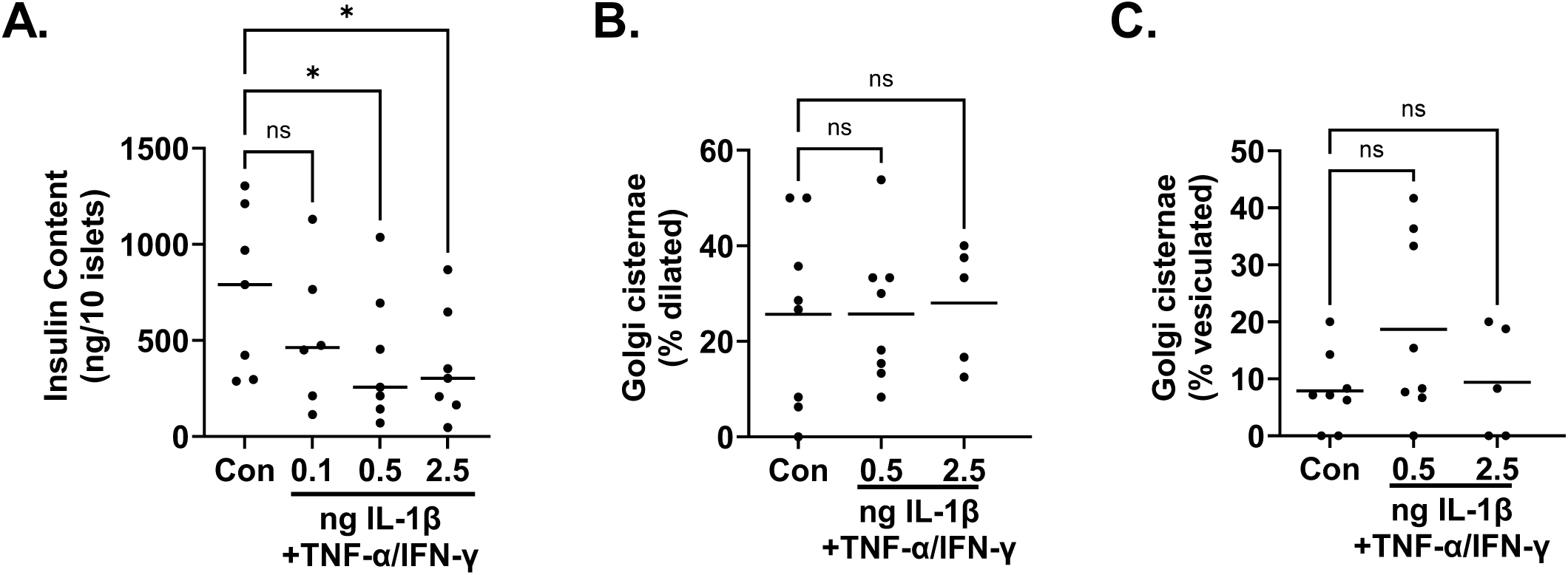
Human insulin content decreases with cytokine treatment, but no change in Golgi morphometrics. Human islets were untreated or treated with cytokine cocktail (IL-1β, 0.1 ng/mL, 0.5 ng/mL, or 2.5 ng/mL; TNF-α, 1000 U/mL; IFN-γ, 750 U/mL) for 48 h. (**A**) Insulin content was determined from whole-cell lysates. Cisternal dilation (**B**) and vesiculation (**C**) was measured from EM images (15 images/donor) as the relative percent of β-cells. Individual donors are presented as data points with the mean as indicated. *p < 0.05 by 1-way ANOVA with repeated measures (**A-C**).

**Supplemental Figure 4.**
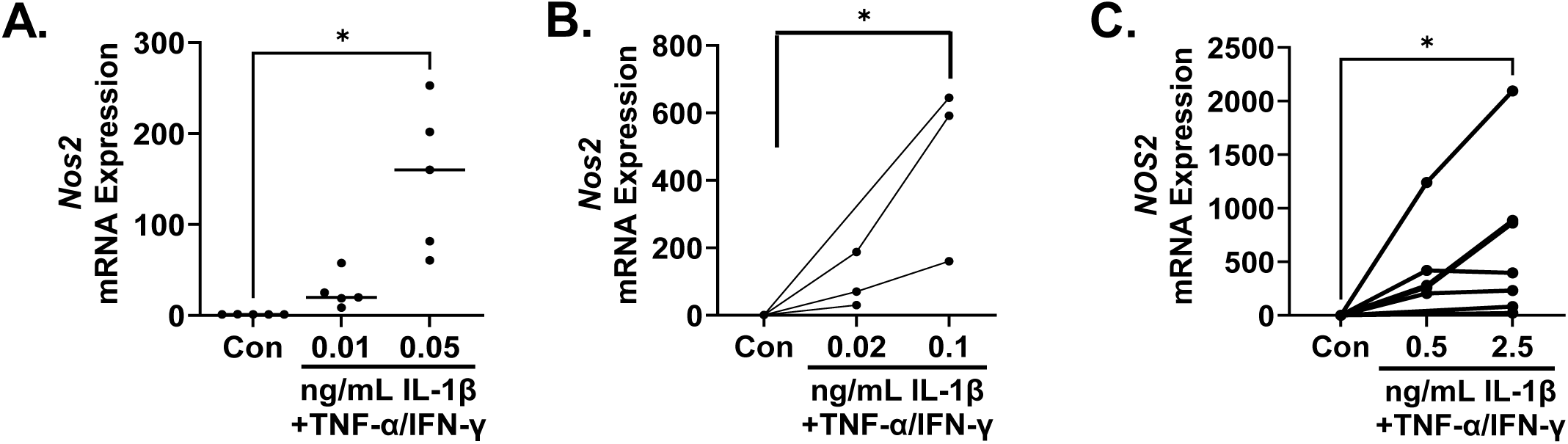
Proinflammatory cytokines induce iNOS expression. (**A**) INS-1 832/3 cells were untreated or treated with cytokine cocktail (IL-1β, 0.01 ng/mL or 0.05 ng/mL; TNF-α, 750 U/mL; IFN-γ, 750 U/mL) for 18 h. (**B**) Mouse islets were untreated or treated with cytokine cocktail (IL-1β, 0.02 ng/mL or 0.1 ng/mL; TNF-α, 750 U/mL; IFN-γ, 750 U/mL) for 18 h. (**C**) Human islets were untreated or treated with cytokine cocktail (IL-1β, 0.5 ng/mL or 2.5 ng/mL; TNF-α, 1000 U/mL; IFN-γ, 750 U/mL) for 24 h. *Nos2* (**A**-**B**) or *NOS2* (**C**) mRNA expression was measured by qRT-PCR. Individual experiments are presented as data points with the mean ± S.D. (**A**) or paired data points from individual mice (**B**) or human donors (**C**). *p < 0.05 by 1-way ANOVA with repeated measures (**A**-**C**).

**Supplemental Figure 5.**
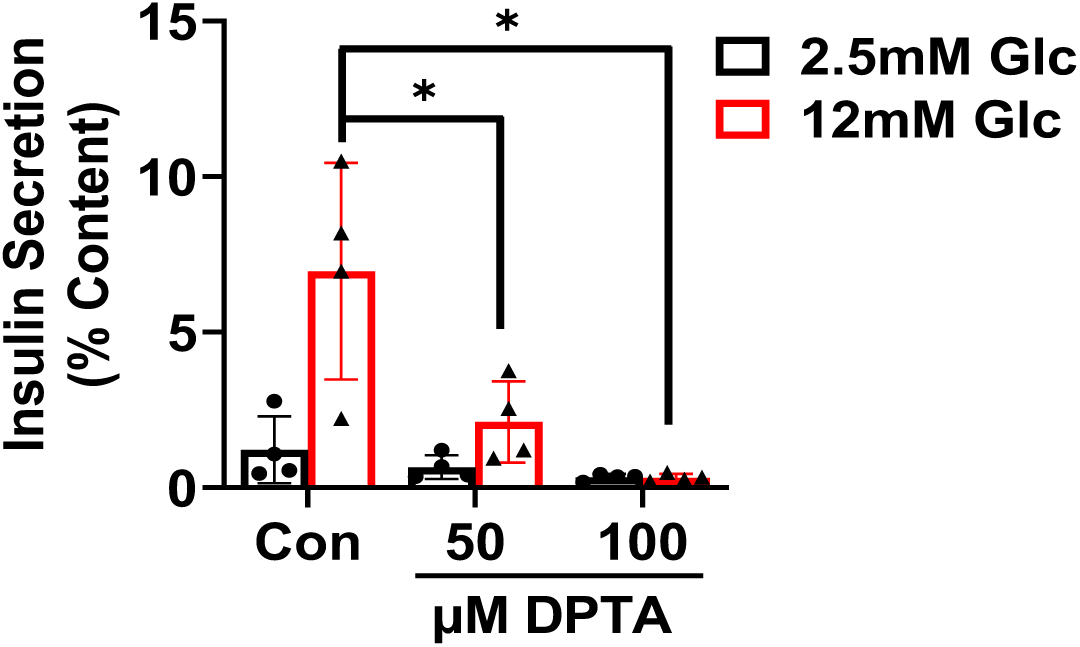
DPTA inhibits insulin secretion. INS1 832/3 cells were untreated or treated with DPTA for 4 h as indicated. Glucose-stimulated insulin secretion was measured by sequential static incubation in media containing 2.5 mM Glc or 12 mM Glc for 1 h each as indicated. Data are normalized to insulin content determined from cell lysates. Individual experiments are presented as data points with the mean ± S.D. *p < 0.05 by 2-way ANOVA with repeated measures.

**Supplemental Figure 6.**
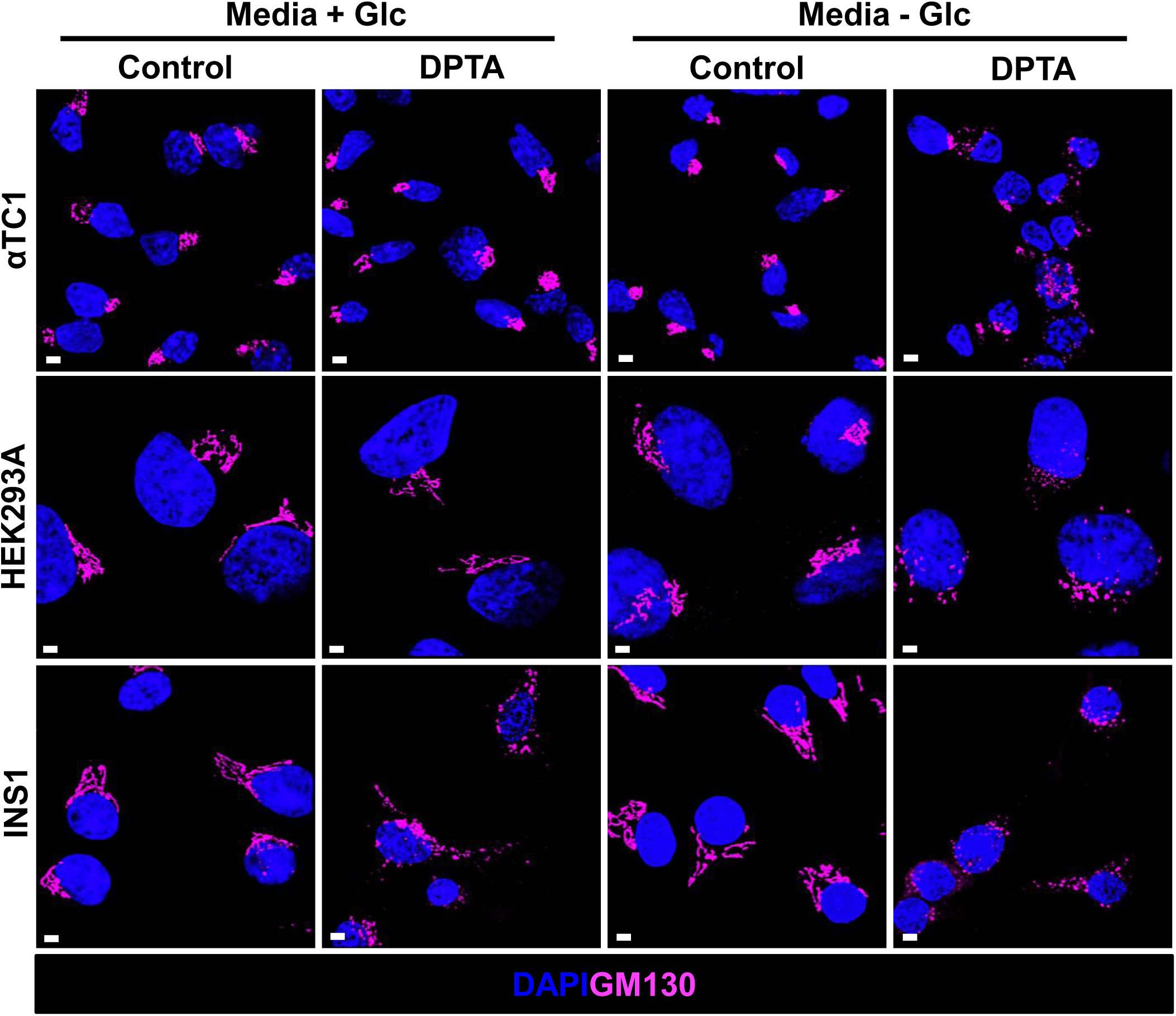
Metabolic differences reveal β-cell specific Golgi responses to NO. αTC1, HEK293A, and INS1 832/3 cells were cultured in growth media with (+) or without (−) glucose and untreated (control) or treated with DPTA (100 μM) for 4 h as indicated. Cells were immunostained for GM130 (magenta) and counterstained with DAPI (blue). Representative images are shown. Scale bar = 3 μm.

**Supplemental Figure 7.**
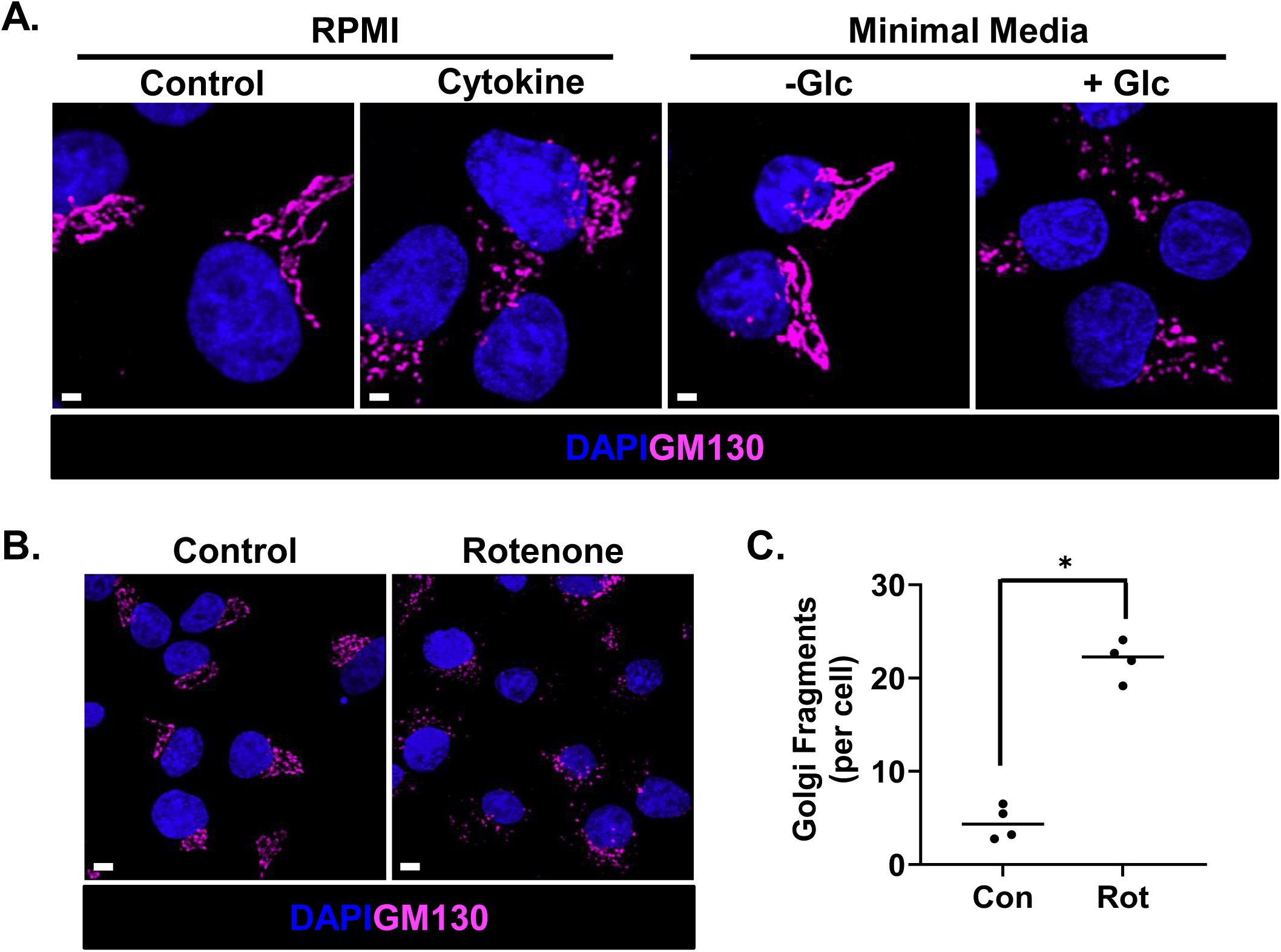
β-cell metabolic activity is required for Golgi structure. (**A**) INS1 832/3 cells were untreated (control) or treated with cytokine cocktail (IL-1β, 0.05 ng/mL; TNF-α, 750 U/mL; IFN-γ, 750 U/mL) for 18 h or cultured in minimal media with (+) or without (−) glucose (Glc, 12 mM) for 8 h. Cells were immunostained for GM130 and counterstained with DAPI. Representative images are shown Scale bar = 3 μm. (**B**-**C**) INS1 832/3 cells were untreated (control) or treated with rotenone (1 µM) for 1 h. Cells were immunostained for GM130 and counterstained with DAPI. (**B**) Representative images are shown. Scale bar = 3 μm. (**C**) Golgi fragments were quantified from GM130 staining per cell. Individual experiments are presented as data points with the mean. *p < 0.05 by Student’s t-test.

## References

1. DiMeglio, L.A., Evans-Molina, C., and Oram, R.A. (2018). Type 1 diabetes. Lancet Lond. Engl. 391, 2449–2462. 10.1016/S0140-6736(18)31320-5.

2. Evans-Molina, C., Sims, E.K., DiMeglio, L.A., Ismail, H.M., Steck, A.K., Palmer, J.P., Krischer, J.P., Geyer, S., Xu, P., and Sosenko, J.M. β Cell dysfunction exists more than 5 years before type 1 diabetes diagnosis. JCI Insight 3, e120877. 10.1172/jci.insight.120877.

3. Broniowska, K.A., Oleson, B.J., and Corbett, J.A. (2014). β-Cell responses to nitric oxide. Vitam. Horm. 95, 299–322. 10.1016/B978-0-12-800174-5.00012-0.

4. Corbett, J.A., Lancaster, J.R., Sweetland, M.A., and McDaniel, M.L. (1991). Interleukin-1 beta-induced formation of EPR-detectable iron-nitrosyl complexes in islets of Langerhans. Role of nitric oxide in interleukin-1 beta-induced inhibition of insulin secretion. J. Biol. Chem. 266, 21351–21354.

5. Corbett, J.A., Wang, J.L., Sweetland, M.A., Lancaster, J.R., and McDaniel, M.L. (1992). Interleukin 1 beta induces the formation of nitric oxide by beta-cells purified from rodent islets of Langerhans. Evidence for the beta-cell as a source and site of action of nitric oxide. J. Clin. Invest. 90, 2384–2391. 10.1172/JCI116129.

6. Southern, C., Schulster, D., and Green, I.C. (1990). Inhibition of insulin secretion by interleukin-1 beta and tumour necrosis factor-alpha via an L-arginine-dependent nitric oxide generating mechanism. FEBS Lett. 276, 42–44. 10.1016/0014-5793(90)80502-a.

7. Welsh, N., Eizirik, D.L., Bendtzen, K., and Sandler, S. (1991). Interleukin-1 beta-induced nitric oxide production in isolated rat pancreatic islets requires gene transcription and may lead to inhibition of the Krebs cycle enzyme aconitase. Endocrinology 129, 3167–3173. 10.1210/endo-129-6-3167.

8. Cardozo, A.K., Ortis, F., Storling, J., Feng, Y.-M., Rasschaert, J., Tonnesen, M., Van Eylen, F., Mandrup-Poulsen, T., Herchuelz, A., and Eizirik, D.L. (2005). Cytokines downregulate the sarcoendoplasmic reticulum pump Ca2+ ATPase 2b and deplete endoplasmic reticulum Ca2+, leading to induction of endoplasmic reticulum stress in pancreatic beta-cells. Diabetes 54, 452–461. 10.2337/diabetes.54.2.452.

9. Corbett, J.A., Sweetland, M.A., Wang, J.L., Lancaster, J.R., and McDaniel, M.L. (1993). Nitric oxide mediates cytokine-induced inhibition of insulin secretion by human islets of Langerhans. Proc. Natl. Acad. Sci. U. S. A. 90, 1731–1735. 10.1073/pnas.90.5.1731.

10. Corbett, J.A., Wang, J.L., Hughes, J.H., Wolf, B.A., Sweetland, M.A., Lancaster, J.R., and McDaniel, M.L. (1992). Nitric oxide and cyclic GMP formation induced by interleukin 1 beta in islets of Langerhans. Evidence for an effector role of nitric oxide in islet dysfunction. Biochem. J. 287 (Pt 1), 229–235. 10.1042/bj2870229.

11. Thomas, H.E., Darwiche, R., Corbett, J.A., and Kay, T.W.H. (2002). Interleukin-1 plus gamma-interferon-induced pancreatic beta-cell dysfunction is mediated by beta-cell nitric oxide production. Diabetes 51, 311–316. 10.2337/diabetes.51.2.311.

12. Scarim, A.L., Heitmeier, M.R., and Corbett, J.A. (1997). Irreversible inhibition of metabolic function and islet destruction after a 36-hour exposure to interleukin-1beta. Endocrinology 138, 5301–5307. 10.1210/endo.138.12.5583.

13. Oleson, B.J., Broniowska, K.A., Naatz, A., Hogg, N., Tarakanova, V.L., and Corbett, J.A. (2016). Nitric Oxide Suppresses β-Cell Apoptosis by Inhibiting the DNA Damage Response. Mol. Cell. Biol. 36, 2067. 10.1128/MCB.00262-16.

14. Stafford, J.D., Shaheen, Z.R., Yeo, C.T., and Corbett, J.A. (2020). Inhibition of mitochondrial oxidative metabolism attenuates EMCV replication and protects β-cells from virally mediated lysis. J. Biol. Chem. 295, 16655–16664. 10.1074/jbc.RA120.014851.

15. Oleson, B.J., and Corbett, J.A. (2018). Dual Role of Nitric Oxide in Regulating the Response of β Cells to DNA Damage. Antioxid. Redox Signal. 29, 1432. 10.1089/ars.2017.7351.

16. Stancill, J.S., Kasmani, M.Y., Khatun, A., Cui, W., and Corbett, J.A. (2022). Cytokine and Nitric Oxide-Dependent Gene Regulation in Islet Endocrine and Nonendocrine Cells. Funct. Oxf. Engl. 3, zqab063. 10.1093/function/zqab063.

17. Stancill, J.S., Kasmani, M.Y., Khatun, A., Cui, W., and Corbett, J.A. (2021). Single-cell RNA sequencing of mouse islets exposed to proinflammatory cytokines. Life Sci. Alliance 4, e202000949. 10.26508/lsa.202000949.

18. Stancill, J.S., Kasmani, M.Y., Cui, W., and Corbett, J.A. (2024). Single Cell RNAseq Analysis of Cytokine-Treated Human Islets: Association of Cellular Stress with Impaired Cytokine Responsiveness. Funct. Oxf. Engl. 5, zqae015. 10.1093/function/zqae015.

19. Stephens, S.B. (2024). Defining Cytokine Responsive and Non-responsive Human β-cells. Function 5, zqae039. 10.1093/function/zqae039.

20. Sims, E.K., Mirmira, R.G., and Evans-Molina, C. (2020). The Role of Beta Cell Dysfunction in Early Type 1 Diabetes. Curr. Opin. Endocrinol. Diabetes Obes. 27, 215–224. 10.1097/MED.0000000000000548.

21. Sims, E.K., Bahnson, H.T., Nyalwidhe, J., Haataja, L., Davis, A.K., Speake, C., DiMeglio, L.A., Blum, J., Morris, M.A., Mirmira, R.G., et al. (2019). Proinsulin Secretion Is a Persistent Feature of Type 1 Diabetes. Diabetes Care 42, 258–264. 10.2337/dc17-2625.

22. Rodriguez-Calvo, T., Chen, Y.-C., Verchere, C.B., Haataja, L., Arvan, P., Leete, P., Richardson, S.J., Morgan, N.G., Qian, W.-J., Pugliese, A., et al. (2021). Altered β-Cell Prohormone Processing and Secretion in Type 1 Diabetes. Diabetes 70, 1038–1050. 10.2337/dbi20-0034.

23. Crawford, S.A., Wiles, T.A., Wenzlau, J.M., Powell, R.L., Barbour, G., Dang, M., Groegler, J., Barra, J.M., Burnette, K.S., Hohenstein, A.C., et al. (2022). Cathepsin D Drives the Formation of Hybrid Insulin Peptides Relevant to the Pathogenesis of Type 1 Diabetes. Diabetes, db220303. 10.2337/db22-0303.

24. Vig, S., Buitinga, M., Rondas, D., Crèvecoeur, I., van Zandvoort, M., Waelkens, E., Eizirik, D.L., Gysemans, C., Baatsen, P., Mathieu, C., et al. (2019). Cytokine-induced translocation of GRP78 to the plasma membrane triggers a pro-apoptotic feedback loop in pancreatic beta cells. Cell Death Dis. 10, 1–13. 10.1038/s41419-019-1518-0.

25. Delong, T., Wiles, T.A., Baker, R.L., Bradley, B., Barbour, G., Reisdorph, R., Armstrong, M., Powell, R.L., Reisdorph, N., Kumar, N., et al. (2016). Pathogenic CD4 T cells in type 1 diabetes recognize epitopes formed by peptide fusion. Science 351, 711–714. 10.1126/science.aad2791.

26. Aaron Wiles, T., Powell, R., Michel, C.R., Scott Beard, K., Hohenstein, A., Bradley, B., Reisdorph, N., Haskins, K., and Delong, T. (2019). Identification of Hybrid Insulin Peptides (HIPs) in Mouse and Human Islets by Mass Spectrometry. J. Proteome Res. 18, 814–825. 10.1021/acs.jproteome.8b00875.

27. Wiles, T.A., Hohenstein, A., Landry, L.G., Dang, M., Powell, R., Guyer, P., James, E.A., Nakayama, M., Haskins, K., Delong, T., et al. (2021). Characterization of Human CD4 T Cells Specific for a C-Peptide/C-Peptide Hybrid Insulin Peptide. Front. Immunol. 12, 668680. 10.3389/fimmu.2021.668680.

28. Cianciaruso, C., Phelps, E.A., Pasquier, M., Hamelin, R., Demurtas, D., Alibashe Ahmed, M., Piemonti, L., Hirosue, S., Swartz, M.A., De Palma, M., et al. (2017). Primary Human and Rat β-Cells Release the Intracellular Autoantigens GAD65, IA-2, and Proinsulin in Exosomes Together With Cytokine-Induced Enhancers of Immunity. Diabetes 66, 460–473. 10.2337/db16-0671.

29. Aguirre, R.S., Kulkarni, A., Becker, M.W., Lei, X., Sarkar, S., Ramanadham, S., Phelps, E.A., Nakayasu, E.S., Sims, E.K., and Mirmira, R.G. (2022). Extracellular vesicles in β cell biology: Role of lipids in vesicle biogenesis, cargo, and intercellular signaling. Mol. Metab. 63, 101545. 10.1016/j.molmet.2022.101545.

30. Casu, A., Nunez Lopez, Y.O., Yu, G., Clifford, C., Bilal, A., Petrilli, A.M., Cornnell, H., Carnero, E.A., Bhatheja, A., Corbin, K.D., et al. (2023). The proteome and phosphoproteome of circulating extracellular vesicle-enriched preparations are associated with characteristic clinical features in type 1 diabetes. Front. Endocrinol. 14, 1219293. 10.3389/fendo.2023.1219293.

31. Lakhter, A.J., Pratt, R.E., Moore, R.E., Doucette, K.K., Maier, B.F., DiMeglio, L.A., and Sims, E.K. (2018). Beta cell extracellular vesicle miR-21-5p cargo is increased in response to inflammatory cytokines and serves as a biomarker of type 1 diabetes. Diabetologia 61, 1124–1134. 10.1007/s00125-018-4559-5.

32. Eizirik, D.L., Cardozo, A.K., and Cnop, M. (2008). The role for endoplasmic reticulum stress in diabetes mellitus. Endocr. Rev. 29, 42–61. 10.1210/er.2007-0015.

33. Pugliese, L.A., De Lorenzi, V., Tesi, M., Marchetti, P., and Cardarelli, F. (2024). Optical Nanoscopy of Cytokine-Induced Structural Alterations of the Endoplasmic Reticulum and Golgi Apparatus in Insulin-Secreting Cells. Int. J. Mol. Sci. 25, 10391. 10.3390/ijms251910391.

34. Bone, R.N., Oyebamiji, O., Talware, S., Selvaraj, S., Krishnan, P., Syed, F., Wu, H., and Evans-Molina, C. (2020). A Computational Approach for Defining a Signature of β-Cell Golgi Stress in Diabetes. Diabetes 69, 2364–2376. 10.2337/db20-0636.

35. Zhang, X., and Wang, Y. (2016). Glycosylation Quality Control by the Golgi Structure. J. Mol. Biol. 428, 3183–3193. 10.1016/j.jmb.2016.02.030.

36. Moremen, K.W., Tiemeyer, M., and Nairn, A.V. (2012). Vertebrate protein glycosylation: diversity, synthesis and function. Nat. Rev. Mol. Cell Biol. 13, 448–462. 10.1038/nrm3383.

37. Xiang, Y., Zhang, X., Nix, D.B., Katoh, T., Aoki, K., Tiemeyer, M., and Wang, Y. (2013). Regulation of protein glycosylation and sorting by the Golgi matrix proteins GRASP55/65. Nat. Commun. 4, 1659. 10.1038/ncomms2669.

38. Bekier, M.E., Wang, L., Li, J., Huang, H., Tang, D., Zhang, X., and Wang, Y. (2017). Knockout of the Golgi stacking proteins GRASP55 and GRASP65 impairs Golgi structure and function. Mol. Biol. Cell 28, 2833–2842. 10.1091/mbc.E17-02-0112.

39. de Graffenried, C.L., and Bertozzi, C.R. (2004). The roles of enzyme localisation and complex formation in glycan assembly within the Golgi apparatus. Curr. Opin. Cell Biol. 16, 356–363. 10.1016/j.ceb.2004.06.007.

40. Ahat, E., Song, Y., Xia, K., Reid, W., Li, J., Bui, S., Zhang, F., Linhardt, R.J., and Wang, Y. (2022). GRASP depletion-mediated Golgi fragmentation impairs glycosaminoglycan synthesis, sulfation, and secretion. Cell. Mol. Life Sci. CMLS 79, 199. 10.1007/s00018-022-04223-3.

41. Luo, Q., Liu, Q., Cheng, H., Wang, J., Zhao, T., Zhang, J., Mu, C., Meng, Y., Chen, L., Zhou, C., et al. (2022). Nondegradable ubiquitinated ATG9A organizes Golgi integrity and dynamics upon stresses. Cell Rep. 40, 111195. 10.1016/j.celrep.2022.111195.

42. Wolfert, M.A., and Boons, G.-J. (2013). Adaptive immune activation: glycosylation does matter. Nat. Chem. Biol. 9, 776–784. 10.1038/nchembio.1403.

43. Ozdilek, A., and Avci, F.Y. (2022). Glycosylation as a key parameter in the design of nucleic acid vaccines. Curr. Opin. Struct. Biol. 73, 102348. 10.1016/j.sbi.2022.102348.

44. Malaker, S.A., Ferracane, M.J., Depontieu, F.R., Zarling, A.L., Shabanowitz, J., Bai, D.L., Topalian, S.L., Engelhard, V.H., and Hunt, D.F. (2017). Identification and Characterization of Complex Glycosylated Peptides Presented by the MHC Class II Processing Pathway in Melanoma. J. Proteome Res. 16, 228–237. 10.1021/acs.jproteome.6b00496.

45. Hohmeier, H.E., Mulder, H., Chen, G., Henkel-Rieger, R., Prentki, M., and Newgard, C.B. (2000). Isolation of INS-1-derived cell lines with robust ATP-sensitive K+ channel-dependent and -independent glucose-stimulated insulin secretion. Diabetes 49, 424–430. 10.2337/diabetes.49.3.424.

46. Zhang, X., Wang, L., Lak, B., Li, J., Jokitalo, E., and Wang, Y. (2018). GRASP55 Senses Glucose Deprivation through O-GlcNAcylation to Promote Autophagosome-Lysosome Fusion. Dev. Cell 45, 245–261.e6. 10.1016/j.devcel.2018.03.023.

47. Bearrows, S.C., Bauchle, C.J., Becker, M., Haldeman, J.M., Swaminathan, S., and Stephens, S.B. (2019). Chromogranin B regulates early-stage insulin granule trafficking from the Golgi in pancreatic islet β-cells. J. Cell Sci. 132, jcs231373. 10.1242/jcs.231373.

48. Stephens, S.B., Edwards, R.J., Sadahiro, M., Lin, W.-J., Jiang, C., Salton, S.R., and Newgard, C.B. (2017). The Prohormone VGF Regulates β Cell Function via Insulin Secretory Granule Biogenesis. Cell Rep. 20, 2480–2489. 10.1016/j.celrep.2017.08.050.

49. Phelps, E.A., Cianciaruso, C., Santo-Domingo, J., Pasquier, M., Galliverti, G., Piemonti, L., Berishvili, E., Burri, O., Wiederkehr, A., Hubbell, J.A., et al. (2017). Advances in pancreatic islet monolayer culture on glass surfaces enable super-resolution microscopy and insights into beta cell ciliogenesis and proliferation. Sci. Rep. 7, 45961. 10.1038/srep45961.

50. Boyer, C.K., Blom, S.E., Machado, Ashleigh E., and Stephens, S.B. (2024). Loss of the Golgi-localized v-ATPase subunit does not alter insulin granule formation or pancreatic islet β-cell function. Am. J. Physiol.-Endocrinol. Metab., (in press).

51. Rabinovitch, A., Suarez, W.L., Thomas, P.D., Strynadka, K., and Simpson, I. (1992). Cytotoxic effects of cytokines on rat islets: evidence for involvement of free radicals and lipid peroxidation. Diabetologia 35, 409–413. 10.1007/BF02342435.

52. McDaniel, M.L., Kwon, G., Hill, J.R., Marshall, C.A., and Corbett, J.A. (1996). Cytokines and nitric oxide in islet inflammation and diabetes. Proc. Soc. Exp. Biol. Med. Soc. Exp. Biol. Med. N. Y. N 211, 24–32. 10.3181/00379727-211-43950d.

53. Sidarala, V., Pearson, G.L., Parekh, V.S., Thompson, B., Christen, L., Gingerich, M.A., Zhu, J., Stromer, T., Ren, J., Reck, E.C., et al. (2020). Mitophagy protects β cells from inflammatory damage in diabetes. JCI Insight 5, e141138, 141138. 10.1172/jci.insight.141138.

54. Oleson, B.J., McGraw, J.A., Broniowska, K.A., Annamalai, M., Chen, J., Bushkofsky, J.R., Davis, D.B., Corbett, J.A., and Mathews, C.E. (2015). Distinct differences in the responses of the human pancreatic β-cell line EndoC-βH1 and human islets to proinflammatory cytokines. Am. J. Physiol. Regul. Integr. Comp. Physiol. 309, R525–534. 10.1152/ajpregu.00544.2014.

55. Subasinghe, W., Syed, I., and Kowluru, A. (2011). Phagocyte-like NADPH oxidase promotes cytokine-induced mitochondrial dysfunction in pancreatic β-cells: evidence for regulation by Rac1. Am. J. Physiol. Regul. Integr. Comp. Physiol. 300, R12–20. 10.1152/ajpregu.00421.2010.

56. Hostens, K., Pavlovic, D., Zambre, Y., Ling, Z., Van Schravendijk, C., Eizirik, D.L., and Pipeleers, D.G. (1999). Exposure of human islets to cytokines can result in disproportionately elevated proinsulin release. J. Clin. Invest. 104, 67–72.

57. Orci, L., Baetens, D., Rufener, C., Amherdt, M., Ravazzola, M., Studer, P., Malaisse-Lagae, F., and Unger, R.H. (1976). Hypertrophy and hyperplasia of somatostatin-containing D-cells in diabetes. Proc. Natl. Acad. Sci. U. S. A. 73, 1338–1342. 10.1073/pnas.73.4.1338.

58. Oleson, B.J., Broniowska, K.A., Yeo, C.T., Flancher, M., Naatz, A., Hogg, N., Tarakanova, V.L., and Corbett, J.A. (2019). The Role of Metabolic Flexibility in the Regulation of the DNA Damage Response by Nitric Oxide. Mol. Cell. Biol. 39, e00153–19. 10.1128/MCB.00153-19.

59. Yeo, C.T., Kropp, E.M., Hansen, P.A., Pereckas, M., Oleson, B.J., Naatz, A., Stancill, J.S., Ross, K.A., Gundry, R.L., and Corbett, J.A. (2023). β-cell-selective inhibition of DNA damage response signaling by nitric oxide is associated with an attenuation in glucose uptake. J. Biol. Chem. 299, 102994. 10.1016/j.jbc.2023.102994.

60. del Valle, M., Robledo, Y., and Sandoval, I.V. (1999). Membrane flow through the Golgi apparatus: specific disassembly of the cis-Golgi network by ATP depletion. J. Cell Sci. 112 (Pt 22), 4017–4029. 10.1242/jcs.112.22.4017.

61. Ranftler, C., Meisslitzer-Ruppitsch, C., Neumüller, J., Ellinger, A., and Pavelka, M. (2017). Golgi apparatus dis- and reorganizations studied with the aid of 2-deoxy-d-glucose and visualized by 3D-electron tomography. Histochem. Cell Biol. 147, 415–438. 10.1007/s00418-016-1515-7.

62. Joshi, G., Chi, Y., Huang, Z., and Wang, Y. (2014). Aβ-induced Golgi fragmentation in Alzheimer’s disease enhances Aβ production. Proc. Natl. Acad. Sci. 111, E1230–E1239. 10.1073/pnas.1320192111.

63. Dal Canto, M.C. (1996). The Golgi apparatus and the pathogenesis of Alzheimer’s disease. Am. J. Pathol. 148, 355–360.

64. Huse, J.T., Liu, K., Pijak, D.S., Carlin, D., Lee, V.M.-Y., and Doms, R.W. (2002). Beta-secretase processing in the trans-Golgi network preferentially generates truncated amyloid species that accumulate in Alzheimer’s disease brain. J. Biol. Chem. 277, 16278–16284. 10.1074/jbc.M111141200.

65. Stieber, A., Mourelatos, Z., and Gonatas, N.K. (1996). In Alzheimer’s disease the Golgi apparatus of a population of neurons without neurofibrillary tangles is fragmented and atrophic. Am. J. Pathol. 148, 415–426.

66. Zhang, X. (2021). Alterations of Golgi Structural Proteins and Glycosylation Defects in Cancer. Front. Cell Dev. Biol. 9, 665289. 10.3389/fcell.2021.665289.

67. Boyer, C.K., Bauchle, C.J., Zhang, J., Wang, Y., and Stephens, S.B. (2023). Synchronized proinsulin trafficking reveals delayed Golgi export accompanies β-cell secretory dysfunction in rodent models of hyperglycemia. Sci. Rep. 13, 5218. 10.1038/s41598-023-32322-z.

68. Ramos-Rodríguez, M., Raurell-Vila, H., Colli, M.L., Alvelos, M.I., Subirana-Granés, M., Juan-Mateu, J., Norris, R., Turatsinze, J.-V., Nakayasu, E.S., Webb-Robertson, B.-J.M., et al. (2019). The impact of proinflammatory cytokines on the β-cell regulatory landscape provides insights into the genetics of type 1 diabetes. Nat. Genet. 51, 1588–1595. 10.1038/s41588-019-0524-6.

69. Pinho, S.S., Alves, I., Gaifem, J., and Rabinovich, G.A. (2023). Immune regulatory networks coordinated by glycans and glycan-binding proteins in autoimmunity and infection. Cell. Mol. Immunol. 20, 1101–1113. 10.1038/s41423-023-01074-1.

70. Kissel, T., Toes, R.E.M., Huizinga, T.W.J., and Wuhrer, M. (2023). Glycobiology of rheumatic diseases. Nat. Rev. Rheumatol. 19, 28–43. 10.1038/s41584-022-00867-4.

71. Ercan, A., Cui, J., Chatterton, D.E.W., Deane, K.D., Hazen, M.M., Brintnell, W., O’Donnell, C.I., Derber, L.A., Weinblatt, M.E., Shadick, N.A., et al. (2010). Aberrant IgG galactosylation precedes disease onset, correlates with disease activity, and is prevalent in autoantibodies in rheumatoid arthritis. Arthritis Rheum. 62, 2239–2248. 10.1002/art.27533.

72. Yang, M.-L., Sodré, F.M.C., Mamula, M.J., and Overbergh, L. (2021). Citrullination and PAD Enzyme Biology in Type 1 Diabetes – Regulators of Inflammation, Autoimmunity, and Pathology. Front. Immunol. 12.

73. Rutter, G.A., Georgiadou, E., Martinez-Sanchez, A., and Pullen, T.J. (2020). Metabolic and functional specialisations of the pancreatic beta cell: gene disallowance, mitochondrial metabolism and intercellular connectivity. Diabetologia 63, 1990–1998. 10.1007/s00125-020-05205-5.

